# The macrocyclizing protease butelase 1 remains auto-catalytic and reveals the structural basis for ligase activity

**DOI:** 10.1101/380295

**Authors:** Amy M. James, Joel Haywood, Julie Leroux, Katarzyna Ignasiak, Alysha G. Elliott, Jason W. Schmidberger, Mark F. Fisher, Samuel G. Nonis, Ricarda Fenske, Charles S. Bond, Joshua S. Mylne

## Abstract

Plant asparaginyl endopeptidases (AEPs) are expressed as inactive zymogens that perform seed storage protein maturation upon cleavage dependent auto-activation in the low pH environment of storage vacuoles. AEPs have attracted attention for their macrocyclization reactions and have been classified as cleavage or ligation specialists. However, we have recently shown that the ability of AEPs to produce either cyclic or acyclic products can be altered by mutations to the active site region, and that several AEPs are capable of macrocyclization given favorable pH conditions. One AEP extracted from *Clitoria ternatea* seeds (butelase 1) is classified as a ligase rather than a protease, presenting an opportunity to test for loss of cleavage activity. Here, making recombinant butelase 1 and rescuing an *Arabidopsis thaliana* mutant lacking AEP, we show butelase 1 retains cleavage functions *in vitro* and *in vivo*. The *in vivo* rescue was incomplete, consistent with some trade-off for butelase 1 specialization toward macrocyclization. Its crystal structure showed an active site with only subtle differences from cleaving AEPs, suggesting the many differences in its peptide binding region are the source of its efficient macrocyclization. All considered, it seems either butelase 1 has not fully specialized or a requirement for auto-catalytic cleavage is an evolutionary constraint upon macrocyclizing AEPs.

## Introduction

Proteases are a large family of enzymes that are responsible for peptide bond breakage and are highly conserved in their mode of catalysis. The residues that comprise the catalytic triad (or dyad) of proteases are highly conserved and an analysis of naturally existing variation among classes of proteases has defined additional geometric constraints on protease active site architecture (Buller and Townsend, 2013).

Asparaginyl endopeptidases (AEPs) are cysteine proteases that cleave peptide bonds following Asn (N) and Asp (D) residues. Like many proteases, AEPs are expressed as zymogens and require auto-activation by cleavage of N- and C- terminal pro-domains at low pH. In plants, AEPs are also known as vacuolar processing enzymes as they process proteins in protein storage vacuoles. Seed storage proteins and AEPs are synthesized on the rough endoplasmic reticulum as precursors before being sorted into separate vesicles. Processing is predicted to occur at the multi-vesicular bodies where both mature and immature seed storage proteins have been shown to co-localize with AEP (Otegui et al., 2006). Ultimately, matured seed storage proteins are trafficked to protein storage vacuoles where they remain until catabolized during germination. An *Arabidopsis thaliana* quadruple *aep* knockout line shows misprocessing of the major seed storage globulins and albumins (Gruis et al., 2004; Kuroyanagi et al., 2005).

In addition to these cleavage functions, AEPs have attracted interest as enzymes that synthesize cyclic peptides (reviewed by (James et al., 2018)). Several structural and molecular studies have helped decipher the structural features necessary for macrocyclization. We have recently shown through structural and biochemical studies of a sunflower AEP that subtle amino acid changes around the active site were able to alter the ratio of cyclized and cleaved reaction products (Haywood et al., 2018) and that all activated AEPs are likely able to perform a macrocyclization reaction at a favorable pH. Mutating a bulky Cys residue located in the substrate channel to a smaller Ala residue greatly increased the rates of macrocyclization activity of an AEP from *Oldenlandia affinis* (Yang et al., 2017). Furthermore, recent sequence analysis and molecular modeling has revealed a range of residues that together contribute to ligation efficiency (Jackson et al., 2018; James et al., 2018; Zauner et al., 2018a).

Given the considerable interest in developing efficient macrocyclizing enzymes as tools for producing highly stable bioactive molecules; butelase 1, the most efficient native cyclizing AEP known to date, has attracted a great deal of attention (Nguyen et al., 2015; Nguyen et al., 2016a; Nguyen et al., 2016b; Bi et al., 2017). However, the practicality of butelase 1 as a tool for synthetic production of cyclic peptides has been limited by an inability to produce it as a recombinant protein (Nguyen et al., 2014).

Although butelase 1 shares high sequence similarity to other AEPs, it exhibits a strong preference for transpeptidation reactions over hydrolysis. Butelase 1 extracted from seeds of the butterfly pea, *Clitoria ternatea* was proposed to have evolved to function as a ligase rather than a protease and would only perform a cleavage reaction in the absence of a suitable nucleophile for macrocyclization (Nguyen et al., 2014). In the presence of a C-terminal tripeptide motif (Asx/His/Val) butelase 1 will accept most N-terminal amino acids for transpeptidation and has been shown to circularize a number of non-native substrates of various sequence compositions and sizes (Nguyen et al., 2014; Nguyen et al., 2015; Hemu et al., 2016). However, when given a fluorogenic substrate recognized and cleaved by AEPs, butelase 1 produced no cleaved product indicating it was incapable of hydrolysis (Nguyen et al., 2014).

With butelase 1 suggested to have evolved to function as an Asx-specific ligase, one might expect it to be unable to self-activate. It is conceivable that butelase 1 is activated by another AEP in *trans* such as butelase 2, which has been shown to have only a cleavage function and to lack cyclizing activity (Serra et al., 2016). To test this hypothesis, we produced recombinant butelase 1 for biochemical studies. We also examined its biological activity *in vivo* by placing a transgene encoding butelase 1 in an *A. thaliana* mutant line that lacks AEP activity. We found recombinant butelase 1 readily self-activated at low pH and would also act as an endopeptidase *in vivo*, partly rescuing the seed protein profile of the *aep* null line. A crystal structure we acquired for butelase 1 showed only subtle differences between its active site and those of AEPs that favor cleavage reactions.

These findings suggest that, although an efficient macrocyclase, butelase 1 either has not yet fully specialized into a macrocyclase or will never fully specialize if the evolutionary constraint of self-processing cannot be overcome in natural systems.

## Results

### Crystal structure of butelase 1

AEPs are expressed as zymogens with a C-terminal pro-domain that ‘caps’ the active site of the core domain. Dissociation of this cap domain occurs upon a shift to a low pH environment where *in trans* self-cleavage at a flexible linker region occurs. Although previous attempts at producing recombinant butelase 1 failed (Nguyen et al., 2014), by using an *E. coli* strain optimized for recombinant proteins with disulfide bonds (Lobstein et al., 2012) we found we could successfully express and purify recombinant butelase 1 as we have done for four other plant AEPs (Bernath-Levin et al., 2015; Haywood et al., 2018). The recombinant protein possessed a 6-His tag at its N-terminus, followed by a Gly-Ser linker and 462 residues of butelase 1 from Ile21 to Val482 (Supplemental Figure 1). This constitutes the full ORF-encoded butelase 1 (482 residues) minus its endoplasmic reticulum signal (20 residues). After purification by immobilized metal affinity chromatography at neutral pH, the purified protein possessed both a small N-terminal pro region and the much larger C-terminal cap that covers the active site.

Crystallization trials of butelase 1 in its inactive form yielded diffraction quality crystals at 3.1 Å resolution. The crystal structure was solved by molecular replacement yielding four molecules in the asymmetric unit each (Figure 1). These molecules were arranged in a manner analogous to that previously reported for *A. thaliana* AEP3, with the asymmetric unit similarly consisting of two dimers formed by interactions between the α6 and α7 helices and stabilized by the C341 specificity loop. Each dimer interaction buries approximately 2500 Å^2^ (Krissinel and Henrick, 2007). The AEP core domain of butelase 1 consists of six β-sheets surrounded by five major α-helices, with several additional short α-helices and β-sheets in the linker regions (Supplemental Figure 2). The C-terminal pro-domain cap, shielding the active site residues Asn59, His165, and Cys207, consists of five helices. The substrate binding site is flanked by two loops between residues Trp236 to Pro245 and Ile289 to Ile293. The latter loop forms an extended, flexible α5-β6 loop, varies in length among various AEPs (Supplemental Figure 2) and only modelled in two molecules of the butelase 1 asymmetric unit due to weak electron density in the other chains.

**Figure 1.**
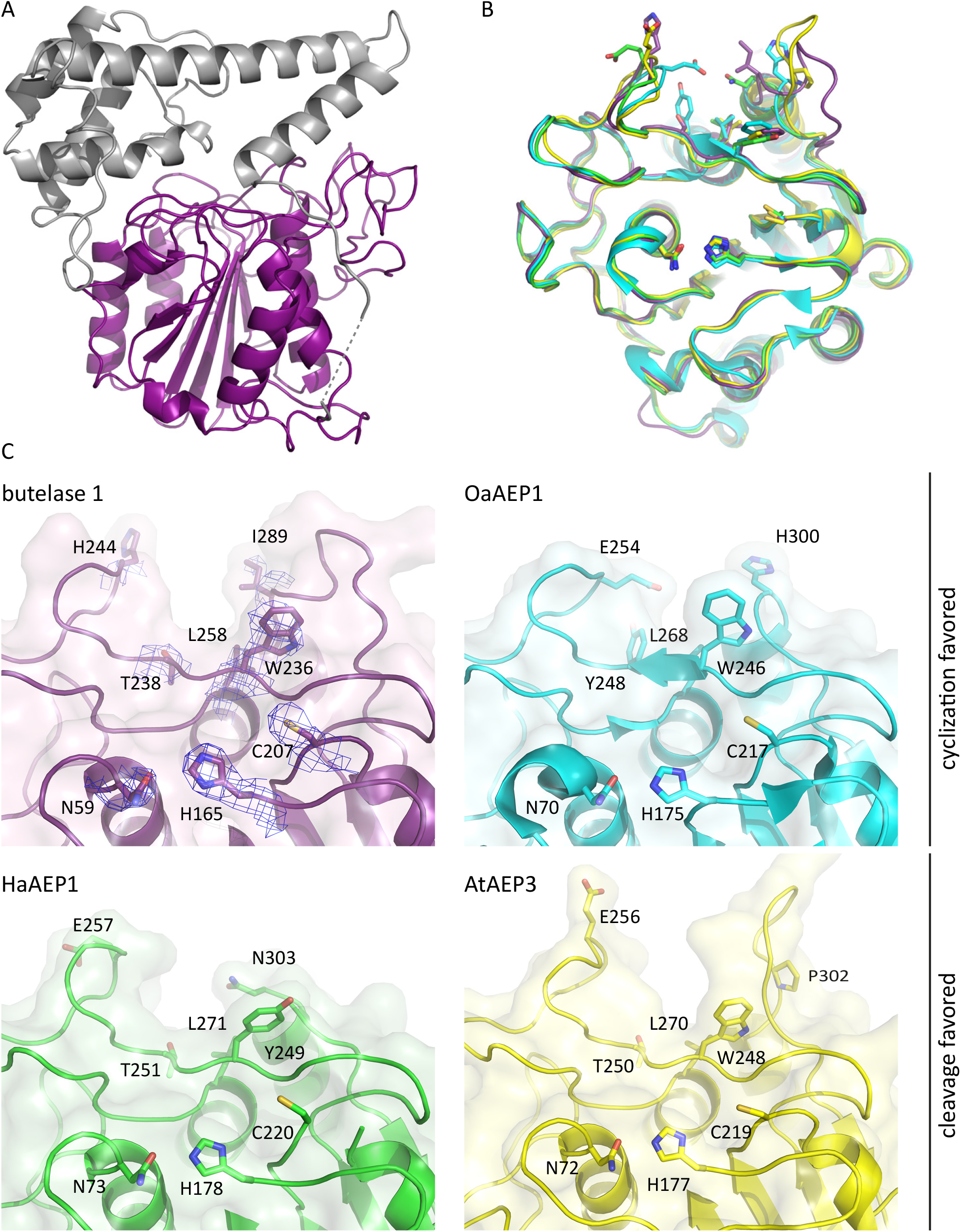
Crystal structure of butelase 1 compared with macrocyclization specialist (OaAEP1) and cleavage-favoring (HaAEP1 and AtAEP3) AEPs. (**A**) Cartoon representation of inactive butelase 1 with core domain and c-terminal cap domain highlighted in purple and gray, respectively. Dashed line represents region of low electron density in linker region between cap and core domain. (**B**) Butelase 1 core domain aligned with OaAEP1, HaAEP1, and AtAEP3 shows high overall structural conservation among AEPs. Catalytic residues and residues in the substrate binding region that may influence catalytic activity shown as sticks. (**C**) Expanded surface and cartoon representations of the predicted catalytic region and substrate binding region highlight subtle changes in residues (shown as sticks) around the substrate binding site that likely contribute to differences in activity. Electron density maps (2 Fobs – Fcalc) for highlighted residues of butelase 1 contoured at 1σ level.

Butelase 1 (Figure 1A) is structurally similar to AEPs from *Oldenlandia affinis* (OaAEP1, PDB ID: 5H0I), the common sunflower *Helianthus annuus* (HaAEP1, PDB ID: 6AZT), and *A. thaliana* (AtAEP3, PDB ID: 5NIJ) with an r.m.s.d. value of 1.1 Å, 0.9 Å, and 1.0 Å over 397, 271, and 422 alpha carbon residues of the most complete chain (B) of butelase 1, respectively (Hasegawa and Holm, 2009). The butelase 1 core domain structure has approximate dimensions of 53 Å × 48 Å × 40 Å and within the core domain the active sites of butelase 1, OaAEP1, HaAEP1, and AtAEP3 show a high degree of structural similarity (Figure 1B). Moreover, comparison between inactive and active forms of AEPs reveal only subtle differences in catalytic residue orientations (Supplemental Figure 3). The catalytic residues in the active site of butelase 1, Asn59, His165, and Cys207, align closely with the catalytic residues of OaAEP1 (Asn70, His175, and Cys217), HaAEP1 (Asn73, His178, and Cys220), and AtAEP3 (Asn72, His177, and Cys219). Furthermore, we chose to model a succinimide (SNN) residue at position 164 due to the preponderance of this aspartimide at this position (N-terminally adjacent to the catalytic His165) in other AEP crystal structures (Dall et al., 2015; Haywood et al., 2018; Zauner et al., 2018a), along with an associated reduction in steric clashes when compared with the modelled alternative Asp164 residue (Supplemental Figure 4). In particular, the short 1.9 Å distance between the OH group of Tyr162 and an Oδ from the side chain of Asp164, which is extended to 2.9 Å with the O5 of SNN, was notable in guiding our decision to model residue 164 as SNN.

The substrate binding sites of butelase 1, on the other hand, showed differences in sequence and structure compared to the aforementioned published plant AEP crystal structures (Figure 1C). All four AEPs have a non-polar Leu residue N-terminal to the start of the α4-helix (Leu258 in butelase 1, Leu268 in OaAEP1, Leu271 in HaAEP1, and Leu270 in AtAEP3) and a polar residue at the start of the C341-loop (Thr238 in butelase 1, Tyr248 in OaAEP1, Thr251 in HaAEP1, and Thr250 in AtAEP3). However, on one side of the entrance to the substrate binding pocket butelase 1 has a bulky Trp residue and a non-polar Ile residue (Trp236 and Ile289). By contrast HaAEP1 has a smaller Tyr residue and a polar Asn residue (Tyr249 and Asn303). OaAEP1 and AtAEP3 both also have a Trp at the former position (Trp246 and Trp248, respectively) but differ in the latter residue with OaAEP1 exhibiting a polar His residue (His300) and AtAEP3 exhibiting a Pro residue (Pro302) that results in a more rigid loop that is shifted away from the binding site. On the opposite side of the entrance to the substrate binding pocket the cyclization specialist butelase 1 has a positively charged residue (His244), whereas the other AEPs possess a negatively charged Glu residue at the same location. Overall, based on these structural differences, the substrate binding region is likely to be responsible for the differences in macrocyclization efficiencies among AEPs.

### Peptide processing by recombinant butelase 1

To be catalytically active, AEPs are thought to undergo auto-catalytic cleavage at an Asx residue *in trans*. However, if the Asx-specific ligase butelase 1 is incapable of cleavage, then it would be unable to self-process. To test this, purified 6-His butelase 1_[21-482]_ with its N-terminal pro-region and C-terminal cap domain was dialyzed at pH 4.0 and found to be capable of autocatalytic cleavage (Figure 2), producing active protein with a mass approximately corresponding to removal of both the N- and C-terminal propeptides (~38 kDa, Figure 2A). This was consistent with the mass of native butelase 1 purified from seeds (Nguyen et al., 2014). This mass was further verified as the active enzyme by incubating the activated protein with a fluorophore-labelled (BODIPY) activity-based probe JOPD1, which is specific for AEP activity (Lu et al., 2015), and then running the mixture on a gel before imaging it (Figure 2B). The ability to self-mature and bind JOPD1 demonstrated that recombinant butelase 1 is capable of performing a cleavage reaction. To provide an opportunity for butelase 1 to perform either a cleavage reaction or its efficient transpeptidation reaction, we incubated purified and self-activated recombinant butelase 1 with an acyclic sunflower trypsin inhibitor (SFTI) synthetic peptide, the seleno-Cys modified SFTI(D14N)-GLDN, which can be processed into both cyclic and acyclic products by other plant AEPs (Bernath-Levin et al., 2015). As predicted, butelase 1 produced cyclic SFTI(D14N) (Figure 2C). In addition to the cyclic product, we also detected a 1626 Da mass consistent with the acyclic-SFTI(D14N) product that would arise from cleavage only (Figure 2C). No mass corresponding to cyclic SFTI(D14N) was detected in a no enzyme control; peaks corresponding to acyclic product were detected in the control that suggested there was some spontaneous cleavage, however these peaks were barely distinguishable above the background (Supplemental Figure 5) and have been observed previously (Bernath-Levin et al., 2015).

**Figure 2.**
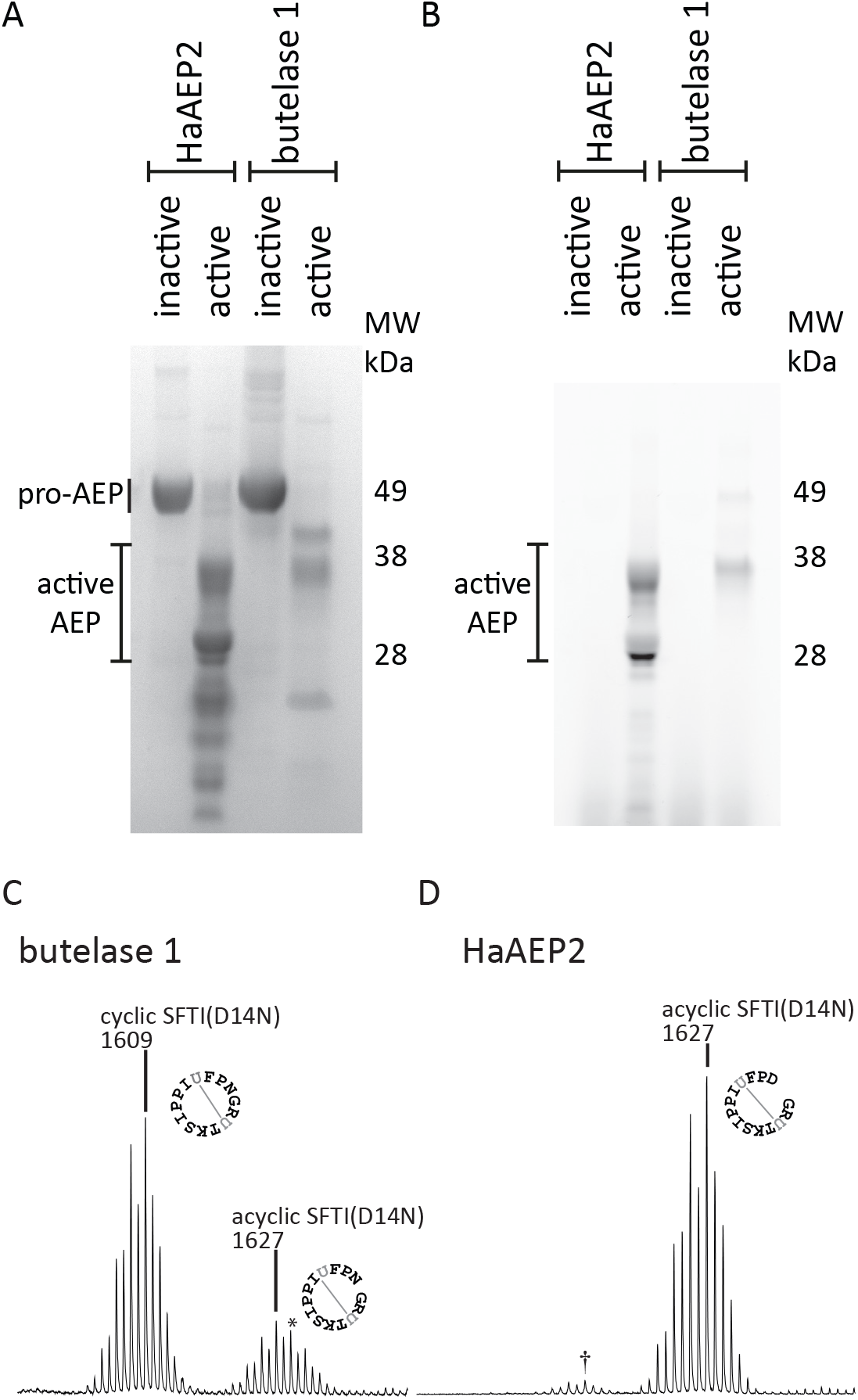
Butelase 1 is capable of performing a cleavage reaction. (**A**) Butelase 1 is capable of auto-activation at low pH. Protein was purified at pH 8.0 and subsequently activated at pH 4.0. The proteins were then dialyzed into activity buffer at pH 5.5 prior to loading onto an SDS-PAGE gel and performing the activity assay. Proteins were visualized with Coomassie stain. The cleavage-favoring HaAEP2 was used as a control illustrating an auto-activation at low pH. (**B**) To confirm which of the masses corresponded to the active protein, protein was incubated with the BODIPY fluorescent probe, JOPD1, which is specific for AEP activity and visualized in SDS-PAGE gel with excitation and emission wavelengths of 532 and 580 nm, respectively. (**C**) Butelase 1 is capable of cleaving a synthetic peptide. Recombinant protein was incubated with synthetic peptide for 24 hours at pH 5.5. Masses consistent with both cyclic SFTI(D14N) and acyclic-SFTI (D14N) were produced following incubation of butelase 1 with the synthetic substrate, consistent with butelase 1 performing both cyclizing and cleaving reactions. A sodium adduct of cyclic SFTI(D14N) resulted in a peak envelope (*) containing both acyclic-SFTI(D14N) and the +22 sodium adduct of cyclic SFTI(D14N). Evidence that these peaks are a result of a sodium adduct is presented in Supplemental Figure 6. (**D**) Recombinant HaAEP2 showed only cleavage ability with the same peptide substrate under the same conditions. The 1611 Da mass (†) is -15 Da from acyclic-SFTI(D14N) so is not cyclic.

As a further control for protease activity we produced recombinant HaAEP2, the second most abundant *AEP* in sunflower seeds based on the number of RNA-seq reads that mapped to sunflower *AEP* transcripts (Bernath-Levin et al., 2015). Only a partial *HaAEP2* sequence was obtained from RNA-seq, however its full-length sequence was obtained by sequencing the cDNA clone that was used to generate an expressed sequence tag found in GenBank (DY926452.1). The construct for recombinant HaAEP2 encoded 440 residues of HaAEP2 with a 6-His and Gly-Ser linker in *lieu* of its ER signal (19 residues) and the predicted N-terminal propeptide (21 residues) (Supplemental Figure 1). HaAEP2, which was purified and activated alongside butelase 1, processed the SFTI(D14N)-GLDN substrate exclusively to a 1626 Da mass with no 1608 Da mass for the cyclic product detectable (Figure 2D). A small amount of a 1611 Da product was observed and this is thought to be due to deamidation of the Asn residue to an acyl group. This mass has been observed previously in studies using the SFTI(D14N)-GLDN substrate (Bernath-Levin et al., 2015). This demonstrated that both butelase 1 and HaAEP2 are capable of performing cleavage reactions. The mass detected for the acyclic-SFTI(D14N) had a mass spectrum that indicated a sodium adduct formed with the cyclic product causing an additional mass 22 Da larger than the protonated form. Sodium adducts of cyclic masses have been observed in previous studies with OaAEP1 (Harris et al., 2015). The presence of the sodium adduct was verified by desalting the sample using solid phase extraction and seeing a reduction, particularly of the 1629 Da peak within the envelope (Supplemental Figure 6). The correct mass was also verified by liquid chromatography-mass spectrometry (LC-MS), which desalted the sample while separating it (Supplemental Figure 6).

The sunflower AEP, HaAEP2, was also shown to be capable of self-maturation using the activity based probe JOPD1; however, it showed heterogeneity in the size of its active form when compared with butelase 1 (Figure 2B).

### Butelase 1 can partially rescue an *aep* null mutant plant

To characterize the activity of butelase 1 *in vivo*, an open reading frame for it under the control of a seed-specific promoter was expressed in an *A. thaliana* mutant lacking endogenous AEP activity. For comparison, the open reading frames of *A. thaliana* AEP2, known be the major seed storage protein processing AEP in *A. thaliana* (Shimada et al., 2003), and *HaAEP2* from sunflower were similarly expressed in the *aep* null line. The seed storage protein profiles of these various transgenic lines were examined by SDS-PAGE and compared to wild type Col-0 or the untransformed *aep* null line (Figure 3). To measure the extent of processing, targeted proteomics was used to quantify the level of tryptic fragments matching either correctly processed or misprocessed seed storage proteins (Figure 4); these values were normalized using tryptic fragments that lack AEP recognition sites, specifically <u>SE</u>ED <u>S</u>TORAGE <u>A</u>LBUMIN 1 (SESA1, At4g27140), SESA3 (At4g27160), and SESA4 (At4g27170), and the globulin CRUCIFERIN3 (CRU3, At4g28520). As expected, the *AtAEP2* transgene fully rescued seed storage protein processing in the *aep* null background (Figure 4B). It is important to note that this full rescue by the *OLEOSIN:AtAEP2* transgene demonstrates that the expression pattern of the *OLEOSIN* promoter is suitable and that the *AEP* gene being intronless does not complicate full and proper rescue by an *AEP* transgene. The heterologous expression of the sunflower AEP2, *HaAEP2*, similarly fully rescued the phenotype of the *aep* null line with no discernible differences in the protein profiles of the HaAEP2 in *aep* lines versus wild type (Figure 4B). The analysis of transgenic HaAEP2 in *aep* was performed twice with comparable results. In transgenic lines where the ORF for butelase 1 was similarly placed under control of the *OLEOSIN* promoter, we observed only partial rescue (Figure 3, Figure 4B). Correctly processed α and β globulins were observed in SDS-PAGE gels; however, several bands corresponding to misprocessed seed storage proteins were seen in extracts from seeds of the *butelase 1* in *aep* line that match bands in untransformed *aep* null (Figure 3).

**Figure 3.**
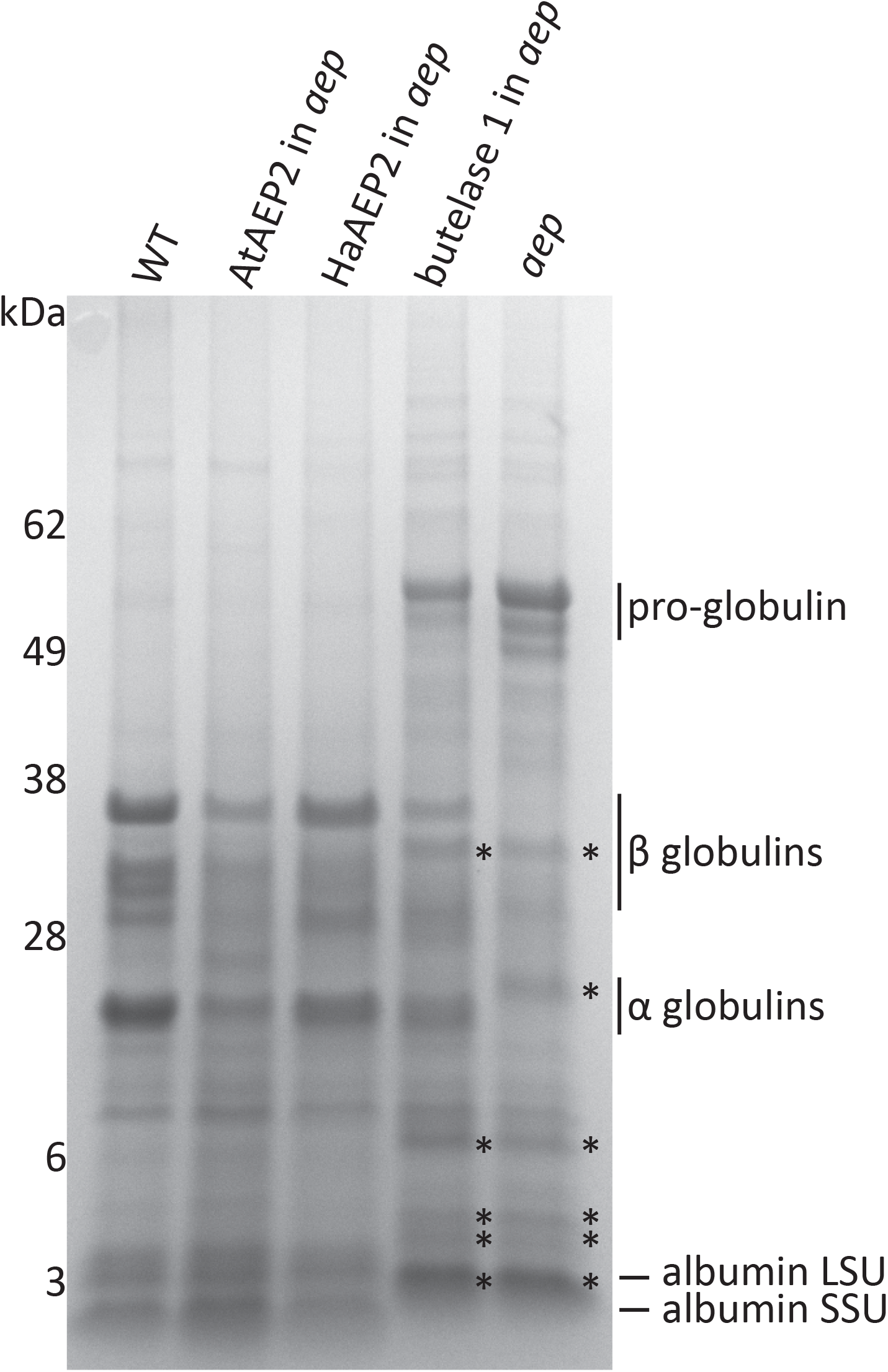
Butelase 1 retains its ability to cleave *in vivo*. Expression of butelase 1 can partially rescue seed storage processing in an *aep* null background. Total protein examined by SDS-PAGE shows expression of *AtAEP2* (lane 2), *HaAEP2* (lane 3), or *butelase 1* (lane 4) under control of the seed-specific *OLEOSIN* promoter can fully or partially rescue the seed protein profile of an *A. thaliana aep* quadruple null mutant (lane 5). Consistent with complete rescue, lanes 1 (Col 0), 2 and 3 have comparable protein profiles indicating correct and complete processing of seed storage proteins by AtAEP2 and HaAEP2. Contrastingly, lane 4 shows an intermediate protein profile between wild type Col-0 (lane 1) and the *aep* quadruple null mutant (lane 5) indicating only partial rescue of the phenotype. Misprocessing events indicated by asterisks can be seen in both protein extracts from *aep* quadruple null mutant seeds and *butelase 1* expressed in *aep*.

**Figure 4.**
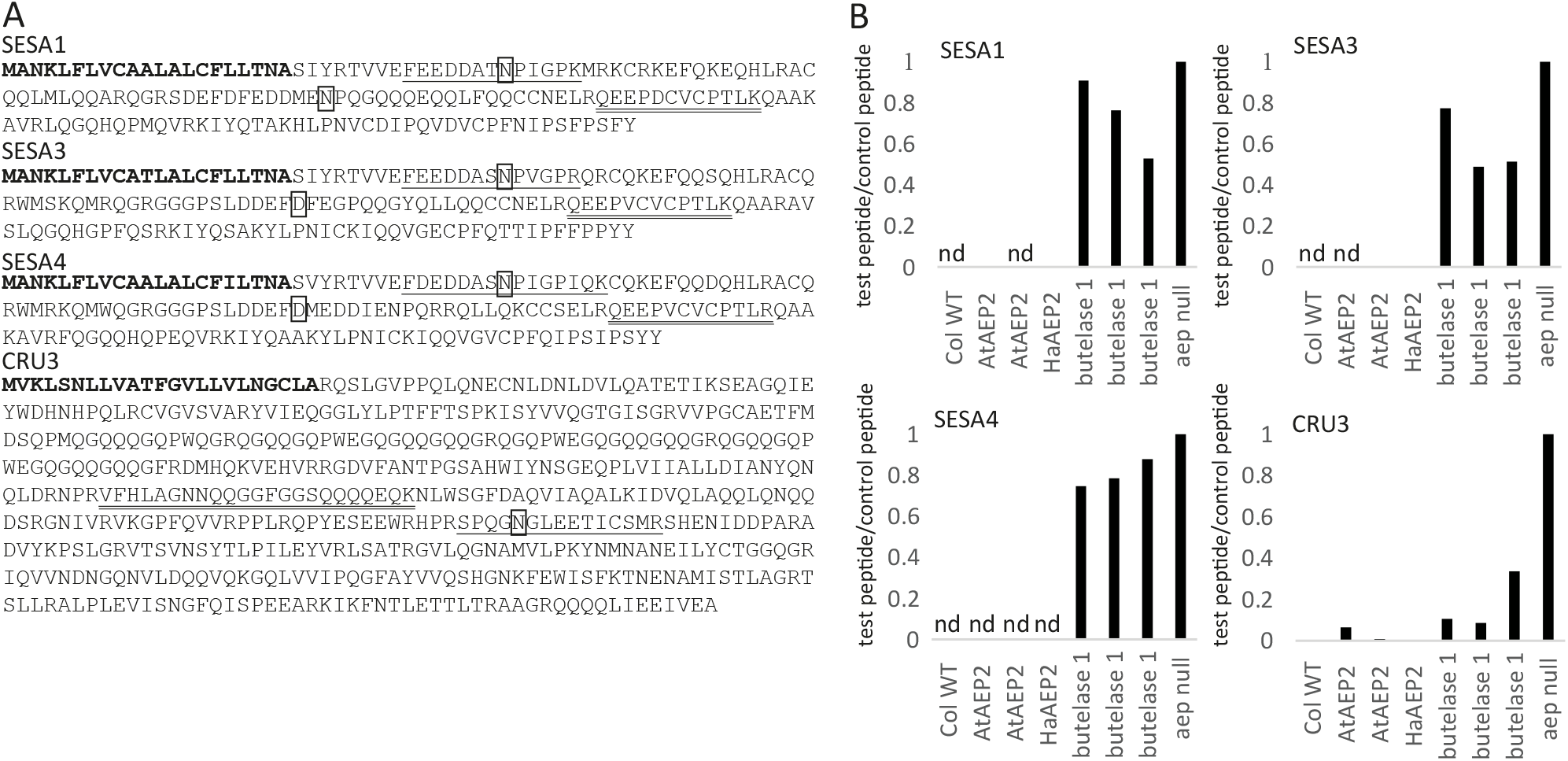
Targeted proteomics monitors *in vivo* activity of several AEPs in an *aep* null. (**A**) Sequences of seed storage proteins used for determining misprocessing events in *aep* mutant plants. ER signal peptides are indicated in bold. The test tryptic fragment with AEP processing site is underlined and the control tryptic fragment lacking AEP processing site is indicated with a double underline. The AEP recognized residues are indicated with black boxes. (**B**) Quantification by multiple reaction monitoring of incorrectly processed seed storage albumins (SESA1, SESA3, SESA4) and globulin (CRU3). Two independent lines with *AtAEP2* and one *HaAEP2* line expressed in an *aep* null background and three independent replicates for *butelase 1* expressed in an *aep* null background are shown. Quantities are given as ratios of test peptide to control peptide and are normalized against *aep* null which is representative of no correct AEP processing events (nd = not detected). Proteomic quantitation of rescue in the *HaAEP2* line was performed in addition to and independently of the data presented in this figure (Table 2, Table 3). **Table**

Gel images were consistent with the quantification provided by targeted proteomics where *AtAEP2* and *HaAEP2* almost fully rescued all of seed storage protein processing in the *aep* null, whereas *butelase 1* rescued 70-90% of CRU3 globulin processing and only 30-40% of albumin processing. (Figure 3, Figure 4,Table 2, Table 3).

**Table 1.** Multiple reaction monitoring transitions used to quantify seed storage protein misprocessing shown in Figure 4B.

| Protein | Peptide Sequence | Precursor<br>m/z | Charge | Product<br>m/z | Charge | Fragment<br>ion |
| --- | --- | --- | --- | --- | --- | --- |
| Test<br>peptide | fragments |  |  |  |  |  |
| SESA1 | FEEDDATNPIGPK | 716.8 | 2 | 726.4 | 1 | y<sub>7</sub><sup>+</sup> |
| SESA3 | FEEDDASNPVGPR | 716.8 | 2 | 525.3 | 1 | y<sub>5</sub><sup>+</sup> |
| SESA4 | FDEDDASNPIGPIQK | 823.4 | 2 | 752.5 | 1 | y<sub>7</sub><sup>+</sup> |
| CRU3 | SPQGNGLEETICSMR | 560.3 | 3 | 553.2 | 1 | y<sub>4</sub><sup>+</sup> |
| Control<br>peptide | fragments |  |  |  |  |  |
| SESA1 | QEEPDCVCPTLK | 738.3 | 2 | 458.3 | 1 | y<sub>4</sub><sup>+</sup> |
| SESA3 | QEEPVCVCPTLK | 730.4 | 2 | 877.4 | 1 | y<sub>7</sub><sup>+</sup> |
| SESA4 | QEEPVCVCPTLR | 744.4 | 2 | 486.3 | 1 | y<sub>4</sub><sup>+</sup> |
| CRU3 | VFHLAGNNQQGGFGGSQQQQEQK | 829.7 | 3 | 660.3 | 1 | y<sub>5</sub><sup>+</sup> |

**Table 2.**
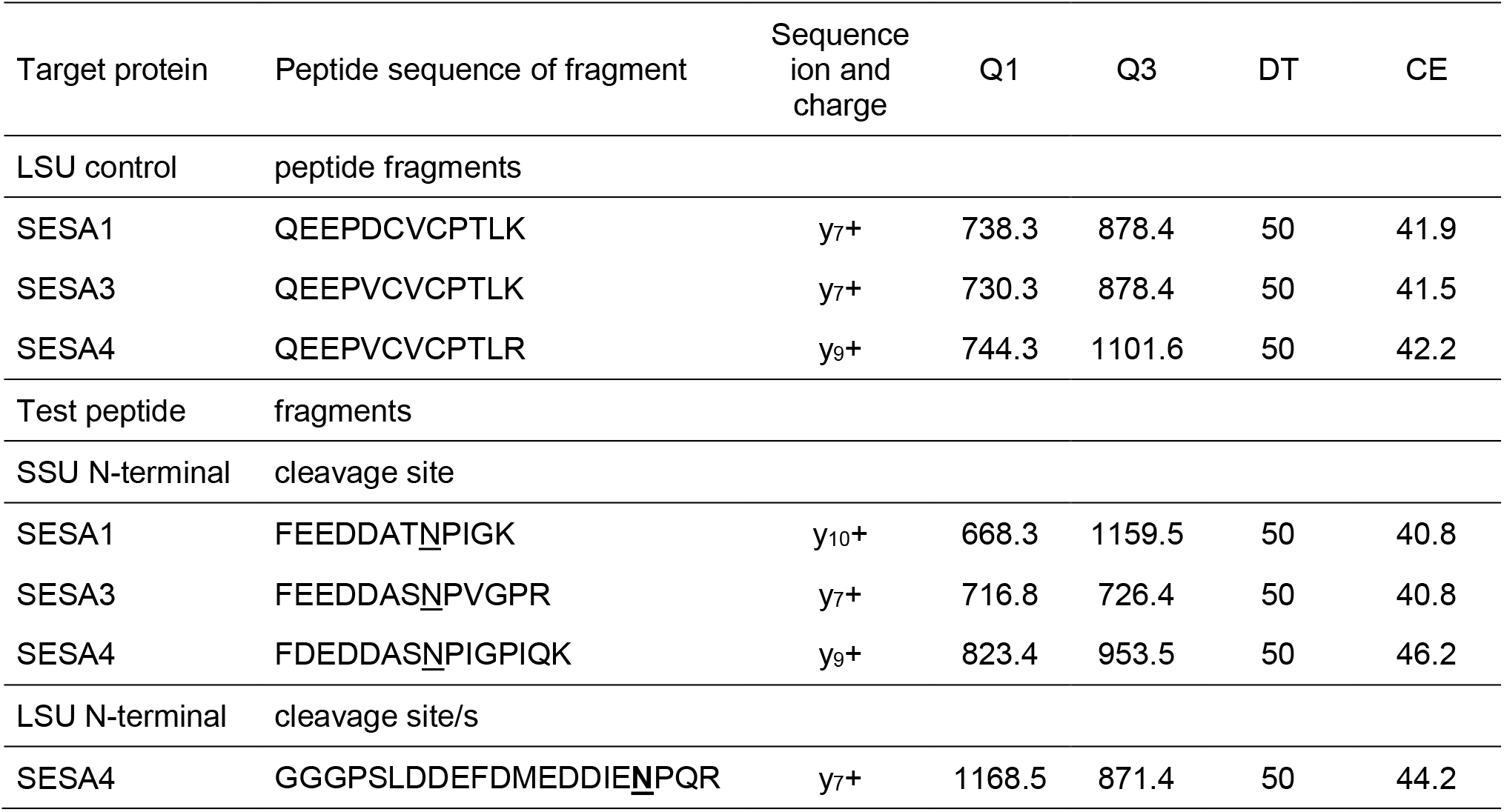
Multiple reaction monitoring transitions used to quantify SEED STORAGE ALBUMIN (SESA) misprocessing shown in Table 3. This analysis was performed independently of the data presented in Figure 4. Multiple reaction monitoring properties include the precursor ion m/z ([M+2H]^2+^) selected in the first quadrupole (Q1), the fragment ion m/z ([M+H]^+^)selected in the third quadrupole (Q3), the dwell time (DT) for each multiple reaction monitoring transition (ms) and the voltage (V) of collision energy (CE). In the test fragments the underlined N (Asn) indicates the *in vivo* cleavage point for AEP. LSU, large subunit. SSU, small subunit.

**Table 3.**
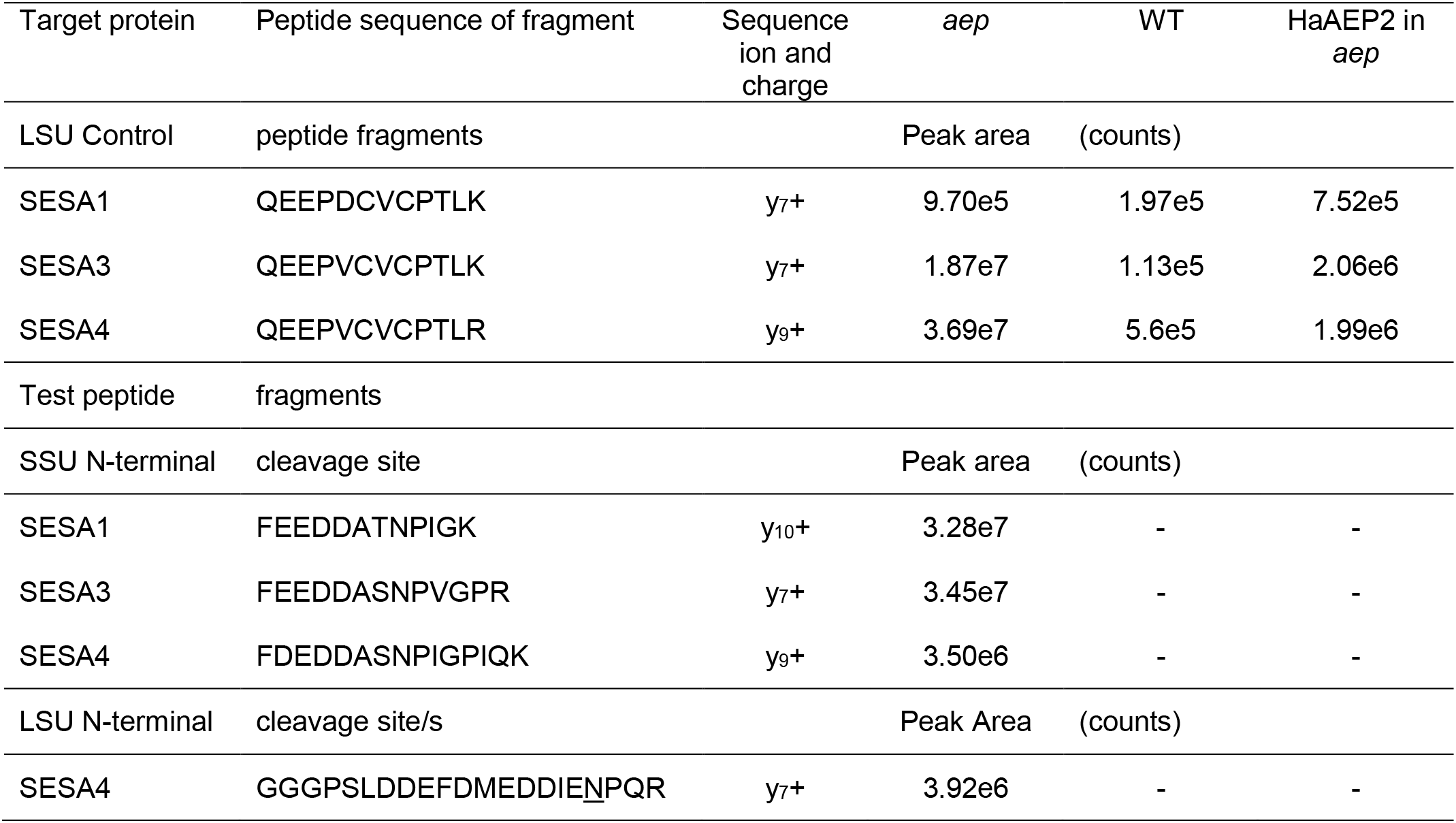
Quantification of SESA tryptic fragments by multiple reaction monitoring. This analysis was performed independently of the data presented in **Figure 4**. The absence of the test SESA fragments in WT and *OLEOSIN:HaAEP2* in *aep* are indicated by a (-). In the test fragments the underlined N (Asn) indicates the *in vivo* cleavage point for AEP. LSU, large subunit. SSU, small subunit.

## Discussion

Several asparaginyl endopeptidases with dual functions, capable of both endopeptidase and transpeptidase reactions, have now been characterized from plants and animals (Bernath-Levin et al., 2015; Dall et al., 2015; Harris et al., 2015). Butelase 1 is one of few examples of asparaginyl endopeptidases that favor transpeptidation over cleavage. The initial study describing butelase 1 activity against non-native substrates found butelase 1 had seemingly a reduced capacity for cleavage activity and so was purported to have evolved to specialize as a ligase (Nguyen et al., 2014). Our study demonstrates that butelase 1 can in fact perform both reactions and will perform a cleavage reaction both *in vitro* and *in vivo*. Using pure, recombinant protein, we demonstrated that butelase 1 was capable of auto-activation, a process that requires cleavage activity. Furthermore, we demonstrated that in the presence of a non-native substrate for macrocyclization, butelase 1 would still perform a cleavage reaction. When expressed in an *A. thaliana* line lacking AEP, a *butelase 1* transgene could partially recover seed storage processing, again an action requiring a cleavage function. Butelase 1 recovered processing of globulins as evidenced by SDS-PAGE analysis and targeted proteomics of seed extracts; however, recovery of albumin processing was minimal with only a partial reduction in the production of misprocessed albumins. Potentially, the endogenous proteases responsible for misprocessing albumins when AEPs are absent are more efficient at cleavage than butelase 1.

The crystal structure of the proenzyme form of butelase 1 bears a strikingly similar structure to those of other AEPs, where an unstructured linker region between the core domain and the cap could provide the flexibility required for self-cleavage of the cap domain upon activation. Furthermore, the crystal structure lends further support for the cap domain forming a dimer interaction surface as suggested previously (Yang et al., 2017; Zauner et al., 2018b). Notably, we have chosen to model a succinimide residue adjacent to the catalytic Cys despite the crystal structure being the pro-form of butelase 1. The electron density at residue 164 is not conclusively illustrative of a SNN residue and quite conceivably the crystal structure could contain a mixture of SNN/Asp at this location. However, we chose to model a SNN residue as this residue is more congruous with the electron density and results in reduced steric clashes. In light of this, our own investigation of the electron density from higher resolution AEP crystal structures 4NOK and 5H0I from inactive mouse and inactive *O. affinis* (Zhao et al., 2014; Yang et al., 2017) indicated that a SNN residue might also be present in these pro-AEP forms (Supplemental Figure 4). Previous observations of SNN residues in the active sites of AEP molecules were made on activated forms of the enzyme which had been subjected to a low pH environment to induce activation. These observations of what appear to be SNN residues in inactive forms of AEP purified at neutral pHs suggests that low pH activation of AEP is not required to form a SNN residue in their active sites.

Recent structural studies of AEPs have identified subtle amino acid changes around the active site and substrate binding domain that could influence their ability to perform a transpeptidation reaction (Yang et al., 2017; Haywood et al., 2018). The crystal structure of butelase 1 shows that residues surrounding the catalytic His are conserved in asparaginyl endopeptidases (Figure 1C). The substrate binding domain shows less conservation between the cyclizing specialists. In the absence of differences between active sites, it is likely that differences in the substrate binding pockets are what define substrate specificity and catalytic efficiency of AEPs, as has been suggested for the improved ligation efficiency of butelase 1 compared to OaAEP1 (Yang et al., 2017). In comparison to previously published crystal structures, butelase 1 exhibits an extended, flexible α5-β6 loop that might become more rigid upon substrate binding and play a role in guiding the N-terminus of a substrate for attack and resolution of a thioacyl intermediate, resulting in peptide cyclization as recently suggested for *A. thaliana* AEP3 (Zauner et al., 2018a). This loop has also recently been highlighted as a ‘marker of ligase activity’ with an extended loop proposed to be absent in ligases, and although butelase 1 does not exhibit such a deletion it was shown to display a more hydrophobic loop than its cleavage-favoring counterpart butelase 2 (Jackson et al., 2018). Moreover, Zauner *et al*. have further substantiated the possible role of a hydrophobic pocket around a conserved Gly residue adjacent to the catalytic His (Gly172 in butelase 1) by molecular dynamics simulations (Zauner et al., 2018a). This region is suspected to play a role in cyclization through the binding of the C-terminal tail of peptide substrates, which is postulated to protect thioacyl intermediates from hydrolysis (Haywood et al., 2018; Zauner et al., 2018a). Further research into these regions with recombinant proteins might discern the subtle amino acid differences that dictate cyclization efficiency and substrate selectivity.

Changes to residues around the substrate binding domain are consistent with results from directed evolution studies of proteins aimed at generating promiscuous functions, which show that mutations leading to novel activity commonly do not impact the protein scaffold or the catalytic residues (Aharoni et al., 2005; Khersonsky et al., 2006). As a result, trade-offs between the new activity and the native activity are minimized (Aharoni et al., 2005). This appears to be the case for dual-function AEPs which do not show any differences in their catalytic residues. Numerous laboratory directed evolution studies have shown that mutations leading to significant improvements in a promiscuous enzyme activity can occur with little effect on the native activity (Aharoni et al., 2005; Khersonsky et al., 2006). However, here we have shown that specializing towards a macrocyclizing activity has resulted in a reduction in cleavage activity for butelase 1 *in vivo* as evidenced by failure to fully recover seed storage processing in the *A. thaliana aep* null mutant.

Evolutionary theory suggests that pre-existing promiscuous enzyme activity is a prerequisite for specialization of novel enzymes following gene duplication and many examples of naturally existing enzymes with promiscuous function exist (Jensen, 1976; Jacob, 1977; Hughes, 1994; Copley, 2014). As we have recently shown, AEPs have an inherent ability to perform ligation under optimal conditions (Haywood et al., 2018). Furthermore, when the SFTI-1 precursor is expressed in wild type *A. thaliana*, a plant lacking cyclic peptides, cyclic SFTI-1 is detectable (Mylne et al., 2011). Therefore, given the high sequence identity of butelase 1 and butelase 2, it is likely that their progenitor sequence possessed both activities.

Although enzymes can possess dual functions, single function enzymes are more common to allow for fine tuning of dosage control and, in cases where there is a negative trade-off for dual functionality, to allow for specialization (Khersonsky et al., 2006). In most cases, gene duplication alleviates the issue of functional constraint as one copy is able to retain the original function and each copy can be regulated independently. This is almost certainly the case for butelase 1 as there is a clear negative trade-off in specializing as a ligase as evidenced by only partial rescue of seed storage processing. However, due to the requirement for auto-catalysis, cleavage ability might be an evolutionary constraint. By maintaining its cleavage ability, butelase 1 can be matured independently from butelase 2. Thus, although gene duplication might have allowed for specialization of butelase 1 into a transpeptidase and butelase 2 to remain an endopeptidase, cleavage activity by butelase 1 might remain so that it can be independently expressed and matured.

Alternatively, there might not be sufficient selection pressure for butelase 1 to completely lose its cleavage function. As observed in directed evolution experiments, without an appropriate selection pressure, complete removal of the original function is difficult to achieve (Tokuriki et al., 2012; Kaltenbach and Tokuriki, 2014). In these cases, mutations leading to reductions of the original activity, followed by optimization of the new activity, corresponded to only weak improvements in the new activity. In nature, the selection pressure for removal of the original activity might not exist once the new activity has been optimized, which is potentially the case for maintenance of the cleavage activity of butelase 1.

In conclusion, this study has demonstrated that butelase 1 has not fully specialized into a cyclizing enzyme. The structural differences between butelase 1 and other AEPs with weak ligation ability suggest that plant AEPs are promiscuous ligases that can readily evolve a macrocyclization capability. This is the first study that has demonstrated successful purification of recombinant butelase 1, whose efficiency makes it useful for the synthetic production of cyclic peptides. Recombinant butelase 1 will expedite future studies as it is now possible to eschew laborious purification of native enzyme from *C. ternatea* seeds (Nguyen et al., 2014; Nguyen et al., 2015; Nguyen et al., 2016b).

Mutagenesis of butelase 1 might enhance its ability to perform macrocyclization reactions and would provide important insights into its evolution.

## Methods

### Recombinant butelase 1 and HaAEP2

A synthetic DNA sequence encoding butelase 1 (Uniprot:A0A060D9Z7), including an N-terminal six-His tag in *lieu* of its ER signal, was designed with codon optimization for *E. coli* (GeneArt) and cloned into pQE30 (Qiagen). Similarly, a synthetic DNA sequence encoding HaAEP2 was designed with codon optimization for *E. coli*, however the sequence included an N-terminal six-His tag in *lieu* of its ER signal, as well as the predicted N-terminal propeptide. The pQE30-HaAEP2 construct was transformed into the T7 SHuffle Express strain of *E. coli* (New England Biolabs) for protein production. The construct encoding butelase 1 was co-transformed into the same *E. coli* strain with the suppressor plasmid pREP4 (Qiagen) for protein production. Proteins were expressed and purified as previously described (Haywood et al., 2018). The proteins were activated by dialysis in activation buffer (20 mM sodium acetate pH 4.0, 5 mM dithiothreitol, 100 mM sodium chloride, 1 mM ethylenediaminetetraacetic acid) for 3 hours at room temperature. Following activation, proteins were dialyzed into activity buffer (0.5 mM dithiothreitol, 100 mM sodium chloride, 1 mM ethylenediaminetetraacetic acid, 20 mM 2-(N-morpholino)ethanesulfonic acid pH 5.5) for 3 hours at room temperature. All proteins were used in assays immediately following dialysis into activity buffer.

### Butelase 1 crystallization and X-ray data collection

Butelase 1 was purified by size-exclusion chromatography (HiLoad 16/600 Superdex 200) in 50 mM Tris (pH 8.0), 50 mM sodium chloride and concentrated to 10-15 mg/mL. Crystal screening was performed using the sitting-drop vapor-diffusion method with 80 μL of reservoir solution in 96-well Intelli-Plates at 20°C. Crystals of butelase 1 were obtained in 10% PEG 8000, 0.2 M sodium chloride, and 0.1 M HEPES (pH 7.5). Single crystals were soaked in mother-liquor supplemented with 20% glycerol as a cryoprotectant prior to being flash-frozen and stored in liquid nitrogen. Data collection was performed at 100 K on the Australian MX1 beamline using a wavelength of 0.9537 Å and diffraction data for crystals were collected to a resolution of 3.1 Å (McPhillips et al., 2002).

### Structure determination and refinement

Diffraction data were processed using XDS and scaled with AIMLESS from the CCP4 program suite (Battye et al., 2011; Winn et al., 2011) in space group P2_1_2_1_2_1_ with unit cell dimensions a = 71.36, b = 147.69, c = 183.33. A sequence alignment of butelase 1 and OaAEP1 was generated using ClustalO and used to create a search model of butelase 1 based on the last common atom of OaAEP1 using CHAINSAW. The structure of butelase 1 was solved by molecular replacement with PHASER using 5H0I as a search model. Manual building and refinement was performed in iterative cycles with Coot and REFMAC5 using the CCP4 program suite. Refinement was performed using jelly-body refinement with sigma 0.02 and non-crystallographic symmetry constraints applied globally. Structural analysis and validation were carried out with Coot and MolProbity (Emsley and Cowtan, 2004; Chen et al., 2010). Crystallographic data and refinement statistics are summarized in (Table 4) with Ramachandran plot values calculated from Coot. Coordinates and structure factors were deposited into the Protein Data Bank (PDB) under accession code 6DHI. Figures illustrating structures were generated using PyMol (Pei et al., 2008; Schrodinger, 2010).

**Table 4.**
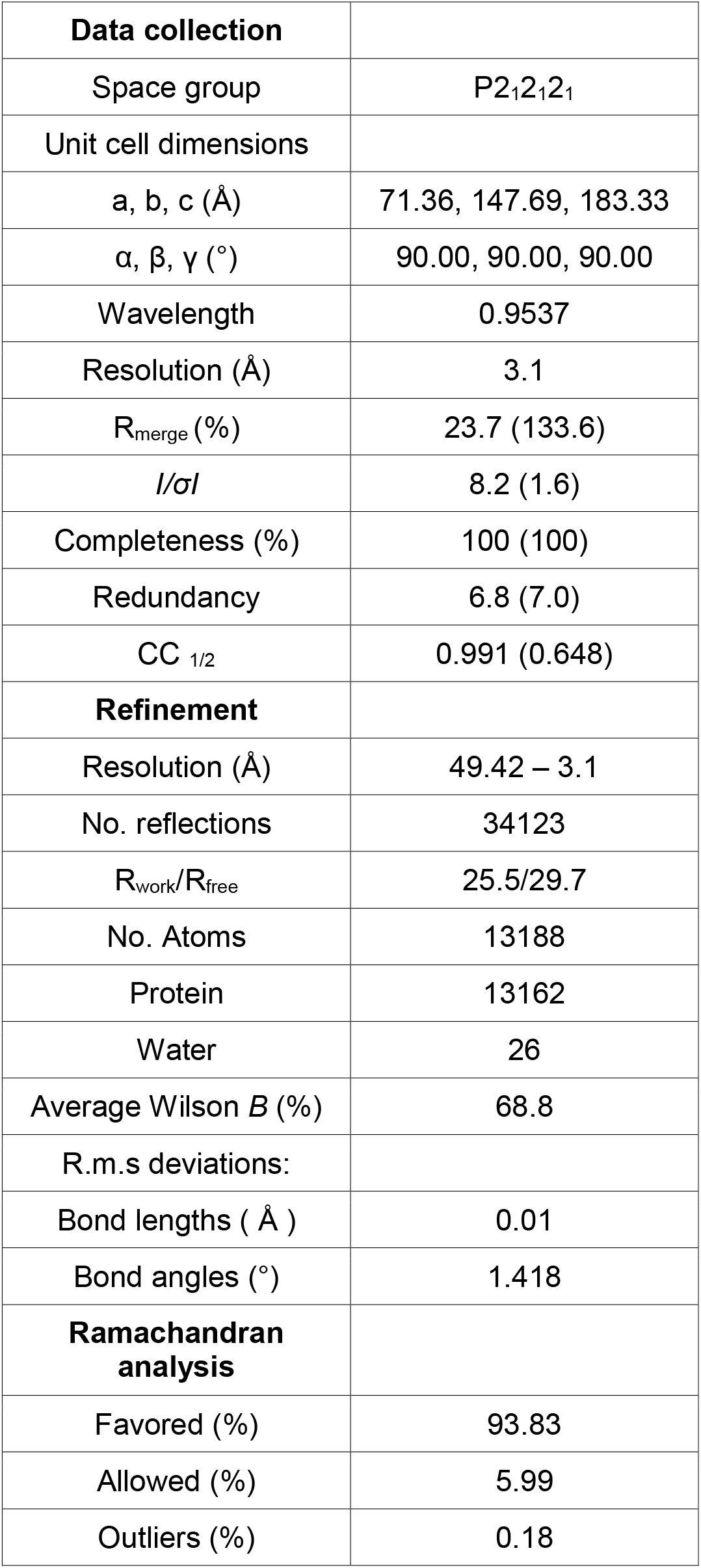
Summary of crystallographic data and refinement statistics. Values in parentheses are for the highest resolution shell.

|  |  |
| --- | --- |
| **Data collection** |  |
| Space group | P2<sub>1</sub>2<sub>1</sub>2<sub>1</sub> |
| Unit cell dimensions |  |
| a, b, c (Å) | 71.36, 147.69, 183.33 |
| α, β, γ (°) | 90.00, 90.00, 90.00 |
| Wavelength | 0.9537 |
| Resolution (Å) | 3.1 |
| R<sub>merge</sub> (%) | 23.7 (133.6) |
| *I*/σ*I* | 8.2 (1.6) |
| Completeness (%) | 100 (100) |
| Redundancy | 6.8 (7.0) |
| CC <sub>1/2</sub> | 0.991 (0.648) |
| **Refinement** |  |
| Resolution (Å) | 49.42 – 3.1 |
| No. reflections | 34123 |
| R<sub>work</sub>/R<sub>free</sub> | 25.5/29.7 |
| No. Atoms | 13188 |
| Protein | 13162 |
| Water | 26 |
| Average Wilson *B* (%) | 68.8 |
| R.m.s deviations: |  |
| Bond lengths ( Å ) | 0.01 |
| Bond angles (°) | 1.418 |
| **Ramachandran analysis** |  |
| Favored (%) | 93.83 |
| Allowed (%) | 5.99 |
| Outliers (%) | 0.18 |

### BODIPY imaging of butelase 1 and HaAEP2

For activity probe analysis 50 μL of AEP at 10 μg/mL was incubated with 0.5 μL of the BODIPY probe JOPD1 at 100 μM (Lu et al., 2015) at room temperature overnight and protected from light. The labelling reaction was stopped by the addition of 10 μL of 6x SDS-PAGE loading buffer (62.5 mM Tris-HCl pH 6.8, 2.5% SDS, 0.002% Bromophenol Blue, 0.7135 M (5%) β-mercaptoethanol, 10% glycerol). Proteins were separated using 4-12% Bis-Tris SDS-PAGE gels as previously described (Lu et al., 2015; Haywood et al., 2018). Labelled proteins were visualized in gel with excitation and emission wavelengths of 532 and 580 nm respectively, using a Typhoon 9500 (GE Healthcare).

### Cleavage and macrocyclization by recombinant AEPs

Both cleavage and macrocyclization activities were determined by incubation of purified recombinant protein with synthetic peptide, seleno-Cys SFTI(D14N)-GLDN. Activated recombinant AEPs at a concentration of 1 mg/mL were incubated with 0.25 mM SFTI(D14N)-GLDN possessing a diselenide bond and 25 μM native (with a disulfide bond) SFTI-1 as an internal standard in activity buffer. Reactions were carried out at 37°C for 24 hours. Synthetic peptides were incubated in the absence of enzyme as a negative control. Most AEPs, along with butelase 1, have shown a preference for Asn hence the use of the SFTI(D14N)-GLDN peptide which possesses an Asn residue in place of the native Asp residue (Nguyen et al., 2014; Bernath-Levin et al., 2015). Products were determined by analysis with an UltraFlex III matrix-assisted laser desorption/time of flight/time of flight (MALDI/TOF/TOF) mass spectrometer (Bruker Daltonics) as described by Bernath-Levin *et al*. (2015).

The presence of a sodium adduct in an unexpectedly wide peak envelope of the acyclic-SFTI(D14N) in the MALDI/TOF/TOF spectra was verified by LC-MS through comparison with the HaAEP2 acyclic product. Reactions were dried and resuspended in 5% acetonitrile 0.1% trifluoroacetic acid and then analyzed using a method described previously (Fisher et al., 2018). Briefly, 2 μL of each sample was injected onto an EASY spray C18 column (75 μm x 150 mm, 3 μm particle size, 10 nm pores; Thermo Scientific) using a Dionex UltiMate 3000 Nano UHPLC system (Thermo Scientific). A 40 min gradient elution was run from 5% acetonitrile to 95% acetonitrile in water with 0.1% formic acid. The UHPLC system was coupled to an Orbitrap Fusion mass spectrometer (Thermo Scientific) for ion detection following electrospray ionization. To further verify the unexpected product, the reaction was desalted using a C18 column and the sample was reanalyzed by MALDI-MS.

### Seed-specific expression of butelase 1, HaAEP2, and AtAEP2 in an *A. thaliana* line lacking AEP

The complete *HaAEP2* open reading frame was amplified by PCR from a sunflower cDNA template using a forward primer JM603 containing an upstream *ClaI* site (5′-tta tcg atA TG AAT CGC CAC CTG CTT ATT CTG-3′) and downstream, a reverse primer JM604 containing a *Sac* I site (5′-atg agc tcG CTG ATT ATA GCA TTA CAT ACA G-3′). AtAEP2 was similarly cloned from *A. thaliana* Columbia (Col-0) cDNA with primers containing the *Cla*I-compatible *Nar*I site upstream and the same *Sac*I restriction enzyme recognition sequence downstream JM384 (5′-aag gcg ccA TGG CTA AGT CTT GCT ATT TCA GAC C-3′) forward; JM385 (5′-aag agc tcA AAA AGT TAG ACG ACC TTA TTG CTA C-3′) reverse. AtAEP2 encodes the AEP primarily responsible for seed storage processing in *A. thaliana* (Shimada et al., 2003; Gruis et al., 2004). A synthetic sequence for *butelase 1* with one silent change (T1071A) to prevent an internal *Sac*I site was designed with a *Cla*I site to the 5’ end and a *Sac*I site to the 3’ end of the sequence. The synthetic *butelase 1* sequence and those for *HaAEP2* and *AtAEP2* were digested with *Cla*I and *Sac*I. The *OLEOSIN* promoter had been cloned previously from Columbia (Col-0) gDNA and was digested with *Xho*I and *Cla*I (Mylne et al., 2011). The *CaMV35S* promoter was removed from pAOV (Mylne and Botella, 1998) by digestion with *Xho*I and *Sac*I. Each *AEP* sequence, the *OLEOSIN* promoter and pAOV were triple ligated as described in Mylne et al. (2011), to generate *Xho*I-*OLEOSIN*-*Cla*I-*AEP*-*Sac*I-*nos* 3’-EcoRI in the pSLJ755I5 binary vector backbone which contains the *bar* gene conferring resistance to glufosinate ammonium herbicide. The binary construct was tri-parental mated into *Agrobacterium tumefaciens* strain LBA4404 and used to transform an *aep* null line essentially as described by Bechtold *et al*. (1993). The *aep* null line contains mutations in all four *AEP* genes, the alleles being *α-1 β-3 γ-1 δ-1* (Kuroyanagi et al., 2005). Seeds from transformed plants (T_0_/T_1_) were sown on soil and sprayed twice with Basta^®^ herbicide at a concentration of 400 mg/L glufosinate ammonium; once after germination when the cotyledons had expanded and then again 3 days later. The T_2_ generation of seeds was collected and sterilized by chlorine gas then sown on MS agar plates containing 80 mg/L glufosinate ammonium. Seeds that showed a 3:1 segregation typical of a single insertion event were collected and sown on soil. Following germination, homozygous plants were selected with several sprays of herbicide. Seeds that showed ~100% herbicide resistance were considered homozygous for a single insertion. Seeds from homozygous plants were used for subsequent analyses. Three independent replicates, representing independent transformation events, for plants expressing *butelase 1* in an *aep* null background and two independent replicates for plants expressing *AtAEP2* in an *aep* null background were analyzed. A single transgenic line was analyzed for HaAEP2 in an *aep* null background.

### Butelase1 maturation of seed storage proteins

Seed storage processing has been shown to be disrupted in the *aep* null background (Gruis et al., 2004). We tested the ability of butelase1, AtAEP2 and HaAEP2 to recover seed storage processing by expressing their encoding genes under the control of the seed specific promoter, *OLEOSIN*, in an *aep* null background. Activity levels were compared to wild type Col-0 and *aep* null Col-0. Approximately 50 mg of seeds were ground in liquid nitrogen with a micro-pestle. To extract protein from seeds, 150 μL of extraction buffer (125 mM Tris-HCl pH 7.0, 7% sodium dodecyl sulfate, 15 mM dithiothreitol, Roche cOmplete™ protease inhibitor [1 tablet/50 mL], 0.5% [w/v] polyvinylpyrrolidone-40) was added to ground tissue and samples were incubated on ice for 20 min on a rocking platform. Soluble protein was separated by centrifuging samples at 10,000 x g for 5 min at 4°C. To precipitate the protein the following steps were taken: to 25 μL of the supernatant (soluble protein fraction), 0.1 mL of chilled methanol was added and the samples were vortexed; then, 25 μL of chilled chloroform was added followed again by vortexing; finally, 75 μL of chilled water was added and the samples were vortexed. Centrifugation at 14,000 x g for 10 minutes at 4°C moved all precipitated protein to the interphase and the aqueous supernatant was discarded. Chilled methanol (0.1 mL) was added and dissolved in the remaining solvent phase, but left the protein interphase as insoluble. To pellet the protein, the sample was mixed by vortexing and centrifuged at 14,000 x g for 10 minutes at 4°C. The supernatant was discarded and the protein pellet was washed three times with chilled acetone.. Following removal of the acetone, the pellet was resuspended in resuspension buffer (7 M urea, 2 M thiourea, 50 mM ammonium bicarbonate, and 10 mM dithiothreitol). Protein was quantified with Bradford reagent and samples were run on an SDS-PAGE gel. For multiple reaction monitoring, samples were alkylated and digested with trypsin as follows: Samples were incubated for 30 minutes at 60°C in 50% volume 100 mM ammonium bicarbonate, 10 mM dithiothreitol. Iodoacetamide was added to a final concentration of 25 mM and the reactions were incubated at room temperature in the dark for one hour. Trypsin was added to the protein samples in a mass ratio of 1:25 and samples were incubated overnight at 37°C. The reaction was stopped with the addition of formic acid to a final concentration of 1% (v/v). Samples were purified by solid phase extraction on Strata™-X 33 μm polymeric reversed phase columns. The columns were conditioned with acetonitrile and equilibrated with 5% (v/v) acetonitrile 0.1% (v/v) formic acid. After loading the columns with the sample, the column was washed once with 1 mL 5% (v/v) acetonitrile 0.1% (v/v) formic acid and once with 1 mL 10% (v/v) acetonitrile 0.1% (v/v) formic acid. The sample was eluted in 0.5 mL 85% (v/v) acetonitrile 0.1% (v/v) formic acid. The eluate was dried under vacuum and resuspended in 5% (v/v) acetonitrile 0.1% (v/v) formic acid to a final concentration of ~1 μg/μL for multiple reaction monitoring. Using an Agilent 1290 Infinity II LC system, 10 μL of each sample were loaded onto an Agilent AdvanceBio Peptide Map column (2.1 × 250 mm, 2.7 μm particle size, P.N. 651750-902), which was heated to 60°C. Peptides were eluted over a 15 minute gradient (0-15 min 3% [v/v] acetonitrile 0.1% [v/v] formic acid to 45% [v/v] acetonitrile 0.1% [v/v] formic acid; 15-15.5 min 45% [v/v] acetonitrile 0.1% [v/v] formic acid to 100% [v/v] acetonitrile 0.1% [v/v] formic acid; 15.5-16 min 100% [v/v] acetonitrile 0.1% [v/v] formic acid to 3% [v/v] acetonitrile 0.1% [v/v] formic acid; 16-30 minutes 3% [v/v] acetonitrile 0.1% [v/v] formic acid) directly into the Agilent 6495 Triple Quadrupole MS for detection. Transitions used for multiple reaction monitoring are given in Table 1.

The following method was used for independent proteomic analysis of seed extracts of *A. thaliana* transgenic lines expressing *HaAEP2* in an *aep* null background compared to wild type and *aep* null seed extracts. Three *A. thaliana* albumins, SESA1, SESA3 and SESA4, were targeted. Trypsin digested protein extracts were analyzed via targeted proteomics. LC-MS/MS data were collected on the AB SCIEX Triple TOF 5600 nanospray coupled to a Shimadzu LC system and the resulting mass spectra were analyzed qualitatively by querying SESA sequences using AB SCIEX Proteinpilot Software 4.0.8085 to identify the individual tryptic fragments. Multiple reaction monitoring was used for quantification of the identified SESA tryptic fragments on the Applied Biosystems 4000 QTRAP nanospray mass spectrometer. Transitions used for multiple reaction monitoring are given in Table 2.

## Accession Numbers

Sequence data from this article can be found in the GenBank/EMBL database under accession MH115430, KF918345, Q39044, AY072508, AY080779, AF446894 and BT002273. Coordinates and structure factors of butelase 1 were deposited into the Protein Data Bank (PDB) under accession code 6DHI.

## Acknowledgements

The authors thank Dan Tawfik for critical insights during project design and Renier van der Hoorn for the BODIPY probe JOPD1. J.S.M. was supported by an Australian Research Council (ARC) Future Fellowship (FT120100013). A.M.J. and S.N. were supported by the Australian Government’s Research Training Program. M.F. was supported by the Australian Government’s Research Training Program and a Bruce and Betty Green Postgraduate Research Scholarship. This research was undertaken in part using the MX1 beamline at the Australian Synchrotron, part of the Australian Nuclear Science and Technology Organisation, and made use of the Australian Cancer Research Foundation detector. J.H. and this work were supported by ARC grant DP160100107 to J.S.M. and Dan Tawfik.

## Author contributions

J.S.M. conceived the study. A.G.E. and J.S.M. cloned *AEP* sequences. A.M.J., J.L. and A.G.E. generated the *A. thaliana* transgenic lines. A.M.J. produced recombinant protein and tested cleavage activity by MALDI analysis. M.F.F. performed LC-MS analysis. A.M.J. and R.F. performed multiple reaction monitoring analysis of seed extracts. K.I. and S.G.N. performed activity based probe assays. A.M.J. and J.H. produced butelase 1 for crystallization. J.H., J.W.S. and C.S.B. solved the crystal structure. A.M.J., J.H., K.I. and J.S.M. wrote the manuscript with contributions from all authors.

## Supplemental Material

**Supplemental Figure 1:**
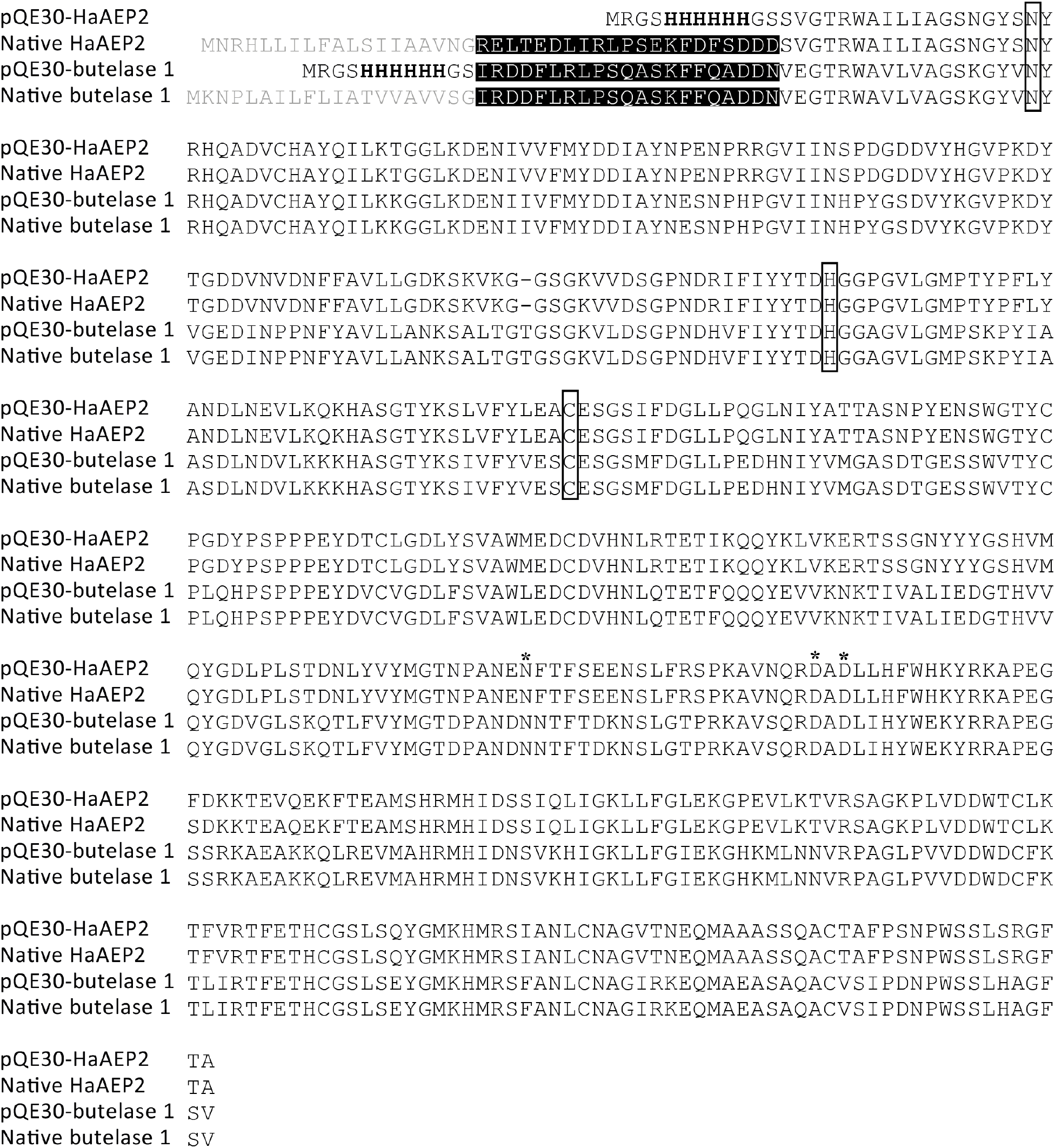
Alignment of the full length native sequences of butelase 1 and HaAEP2 and the synthetic sequences designed for generation of recombinant protein. ER-signals are indicated in gray. N-terminal propeptides are indicated in white font with black highlight. The six-His tag are indicated in bold. Asterisks indicate sites of auto-cleavage. The conserved residues of the catalytic triad are indicated with black boxes.

**Supplemental Figure 2:**
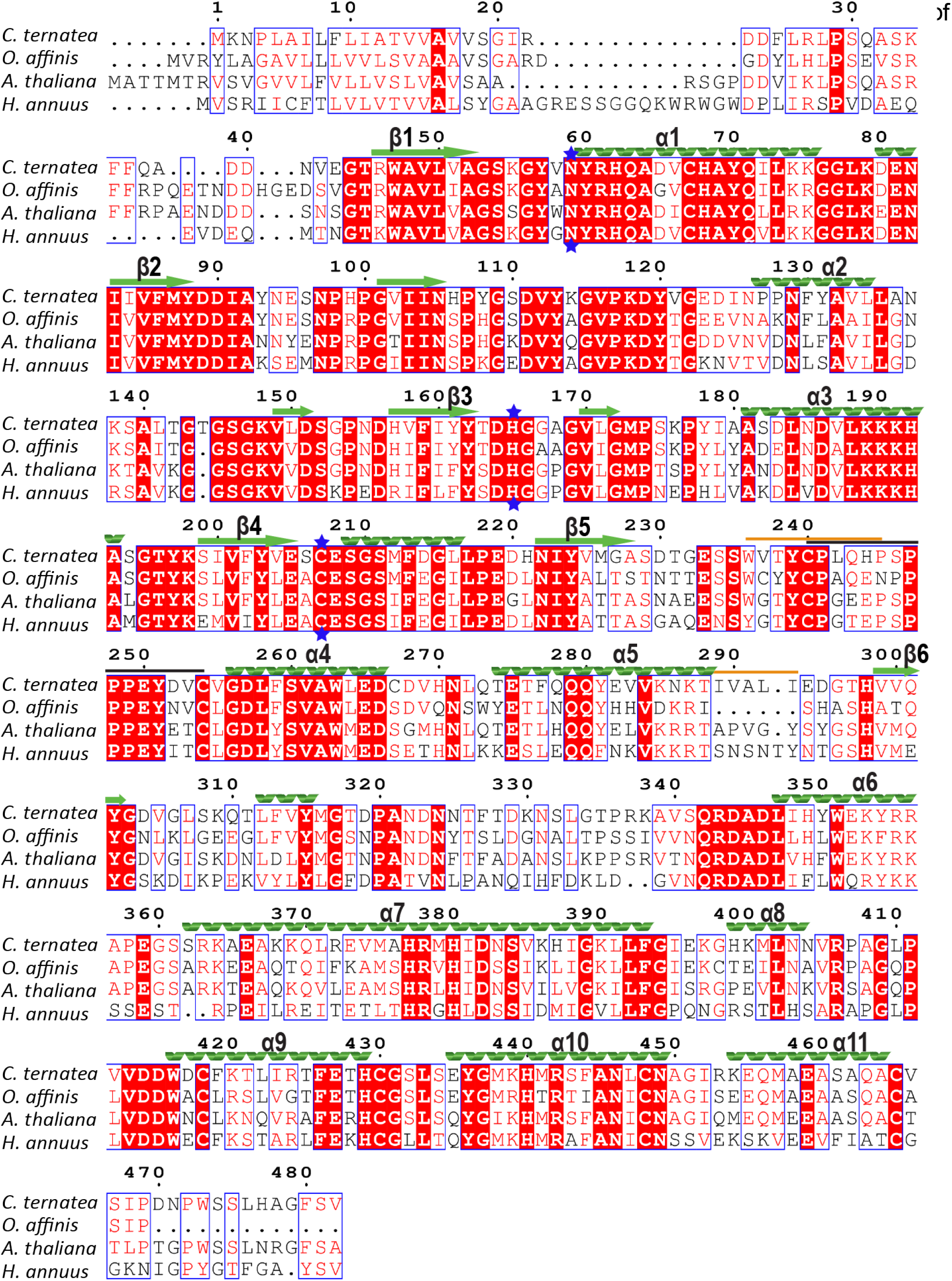
AEPs for which protein structure is known. Sequence alignment generated with PDB files and sequences of AEPs from *C. ternatea* butelase 1 (Uniprot:A0A060D9Z7 PDB: 6DHI), *O. affinis* (Uniprot: A0A0N9JZ32 PDB: 5H0I) *A. thaliana* (Uniprot: Q39119 PDB: 5NIJ) *H. annuus* (Uniprot: A0A0G2RI59 PDB: 6AZT). Identical residues shown with white text and red background, similar residues shown with red text. Conserved secondary structure β- sheets and α-helices shown above sequences with green arrows and green helices, respectively, and with major β-sheets and α-helices labelled. Catalytic triad residues highlighted with blue stars. C341-loop region highlighted with black bar above sequence. Regions of amino acid diversity in substrate binding site highlighted with orange bar above sequence (Pei et al., 2008).

**Supplemental Figure 3:**
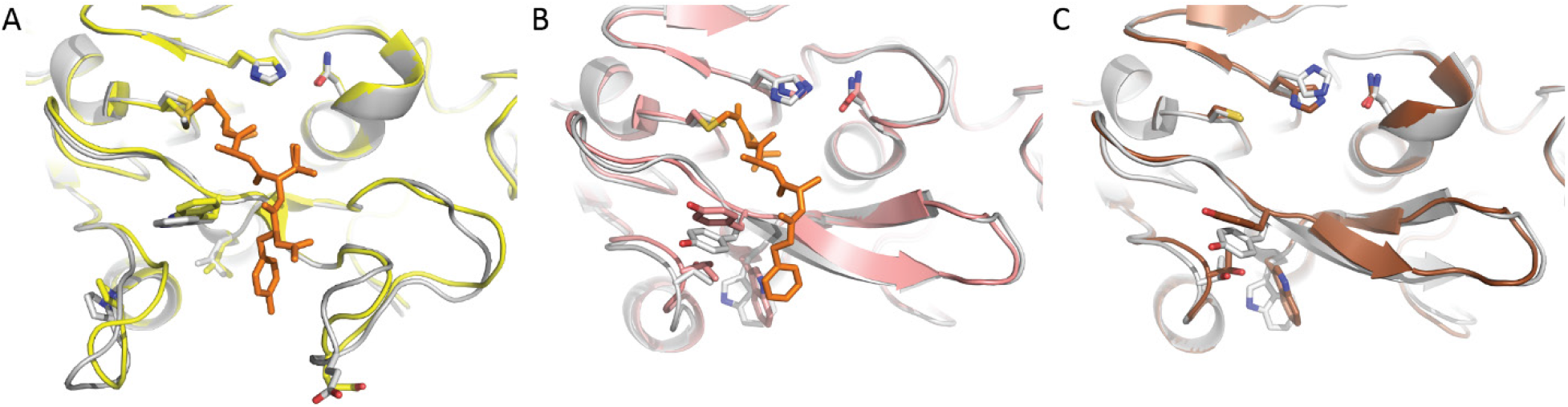
Comparison of available crystal structures of active and inactive forms of AEPs catalytic core domain. Cartoon representation of core domains with residues thought to play a role in catalytic activity shown as sticks. (**A**) *A. thaliana* AEP3, active yellow with bound chloromethylketone inhibitor shown as orange sticks (PDB: 5OBT), inactive gray (PDB: 5NIJ). (**B**) Human AEP, active pink with bound chloromethylketone inhibitor shown as orange sticks (PDB: 4AWB), inactive gray (PDB: 4FGU). (**C**) Mouse AEP, active brown (PDB: 4NOJ), inactive gray (PDB: 4NOK).

**Supplemental Figure 4:**
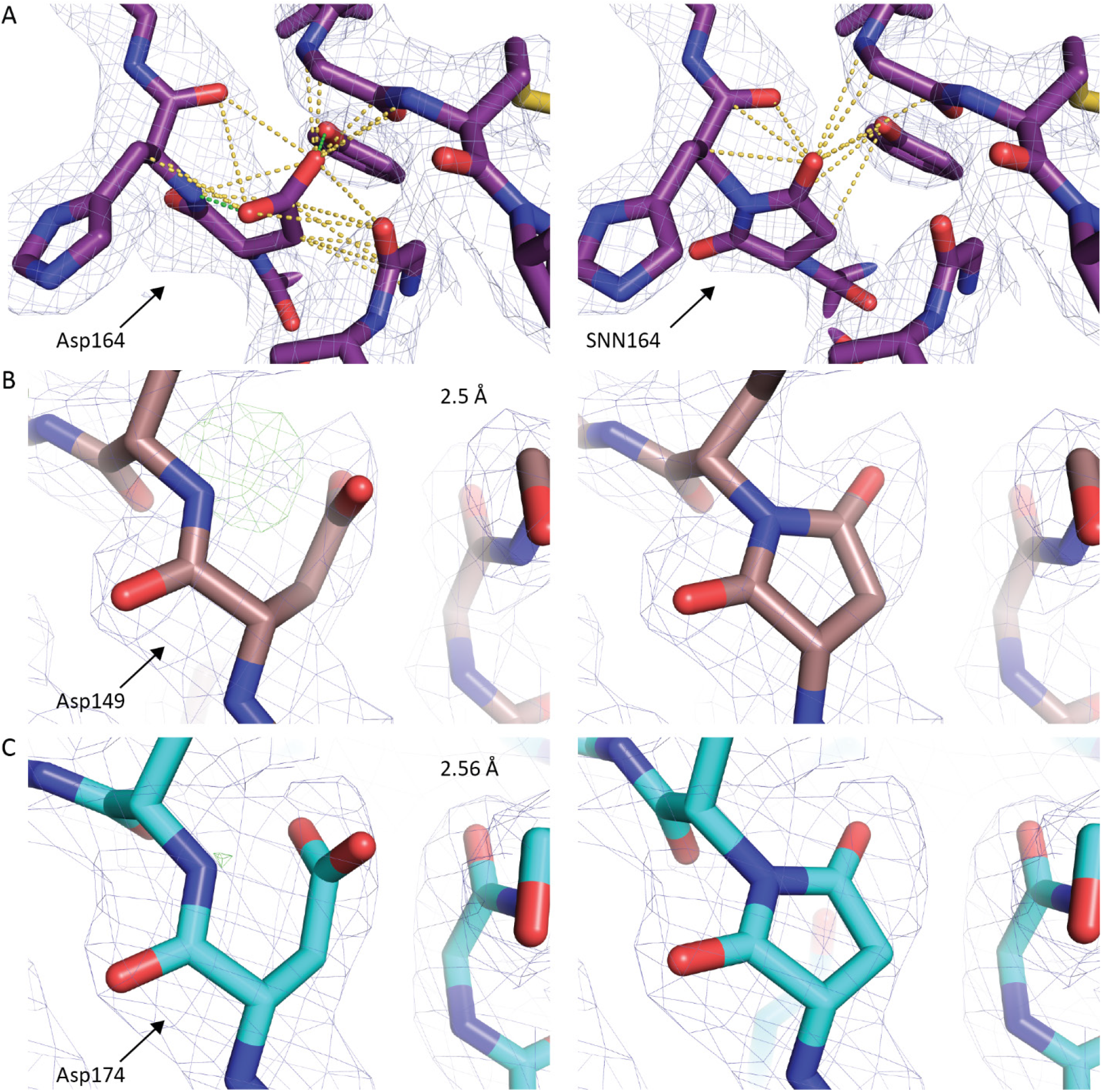
Asp residues modelled as SNN. (**A**) Comparison of alternative models for butelase 1 residue 164. Asp164 was chosen to be modelled as a succinimide residue (SNN) in all four chains of the butelase 1 PDB as the modelled SNN residue was more congruous with the electron density and showed reduced steric clashes when compared with the modelled Asp164 residue. Electron density maps (2 F_obs_ – F_calc_) contoured at 1 σ level. Contacts to the side chain of residue 164 of ≤ 3.5 Å and ≤ 2.0 Å illustrated with yellow and green dashed lines, respectively. Also shown are examinations of electron density for the equivalent Asp residue (left image) in higher resolution structures for inactive AEPs from (**B**) mouse (PDB:4NOK) and (**C**) *O. affinis* (PDB: 5H0I) with additional omit maps (F_obs_ – F_calc_) contoured at 3 σ level (green). Models of SNN residues are presented in the images on the right.

**Supplemental Figure 5:**
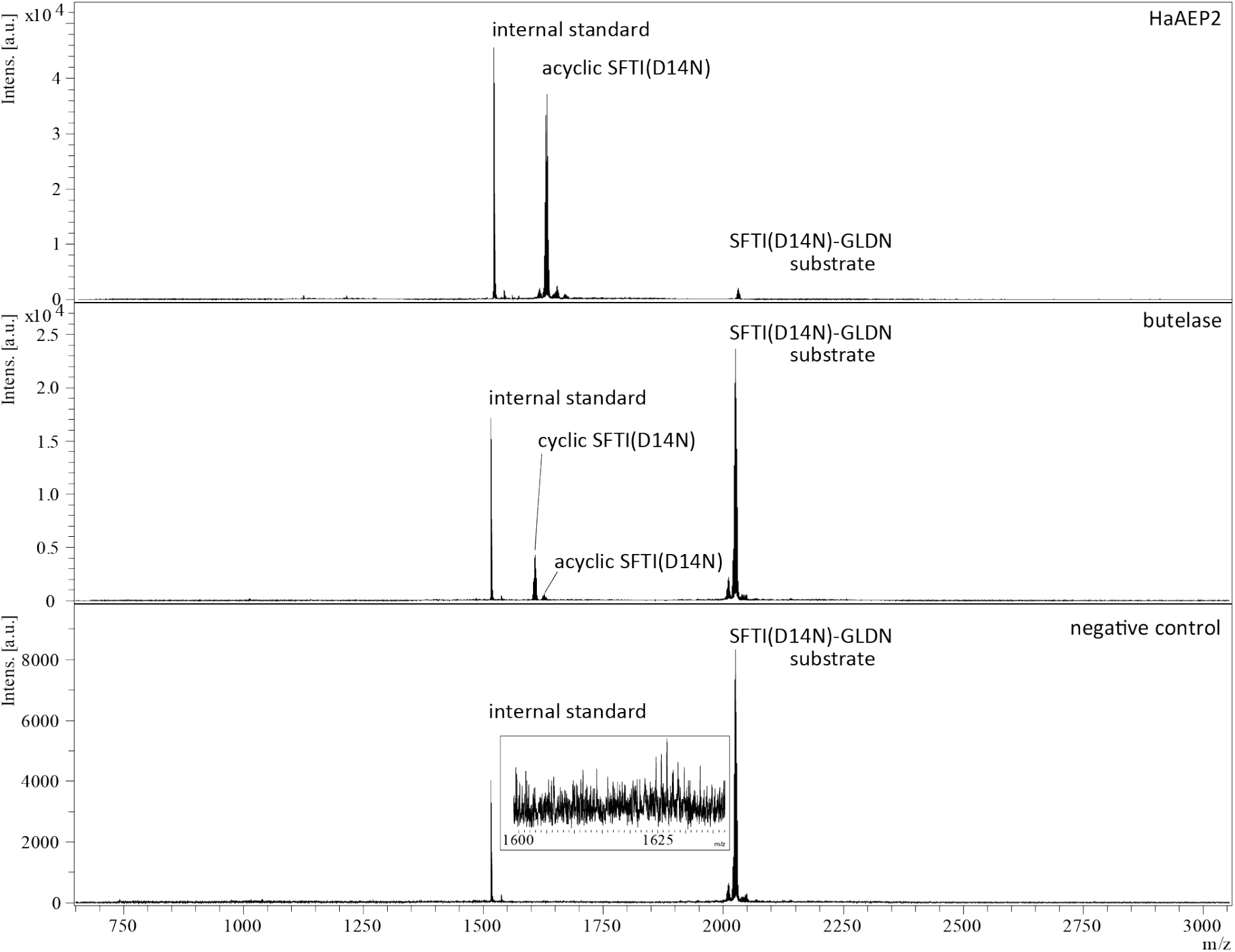
MALDI/TOF spectra of processing of SFTI(D14N)-GLDN by HaAEP2 and butelase 1. SFTI(D14N)-GLDN was incubated under the same conditions without any enzyme present as a negative control. In each condition a mass of 2027 Da corresponding to the substrate, seleno-Cys modified SFTI(D14N)-GLDN, is detected indicating incomplete processing after 24 hours. A mass of 1626 Da corresponding to acyclic SFTI(D14N) and a mass of 1608 Da corresponding to cyclic SFTI(D14N) were detected when the substrate was incubated with butelase 1, whereas only a mass corresponding to acyclic SFTI(D14N) was detected when the substrate was incubated with HaAEP2. No mass corresponding to cyclic SFTI(D14N) was detected in the negative control. A weak signal could be detected for acyclic SFTI(D14N) in the negative control, but was negligible compared to the test samples. Native cyclic SFTI was included as an internal control with a mass of 1515 Da.

**Supplemental Figure 6:**
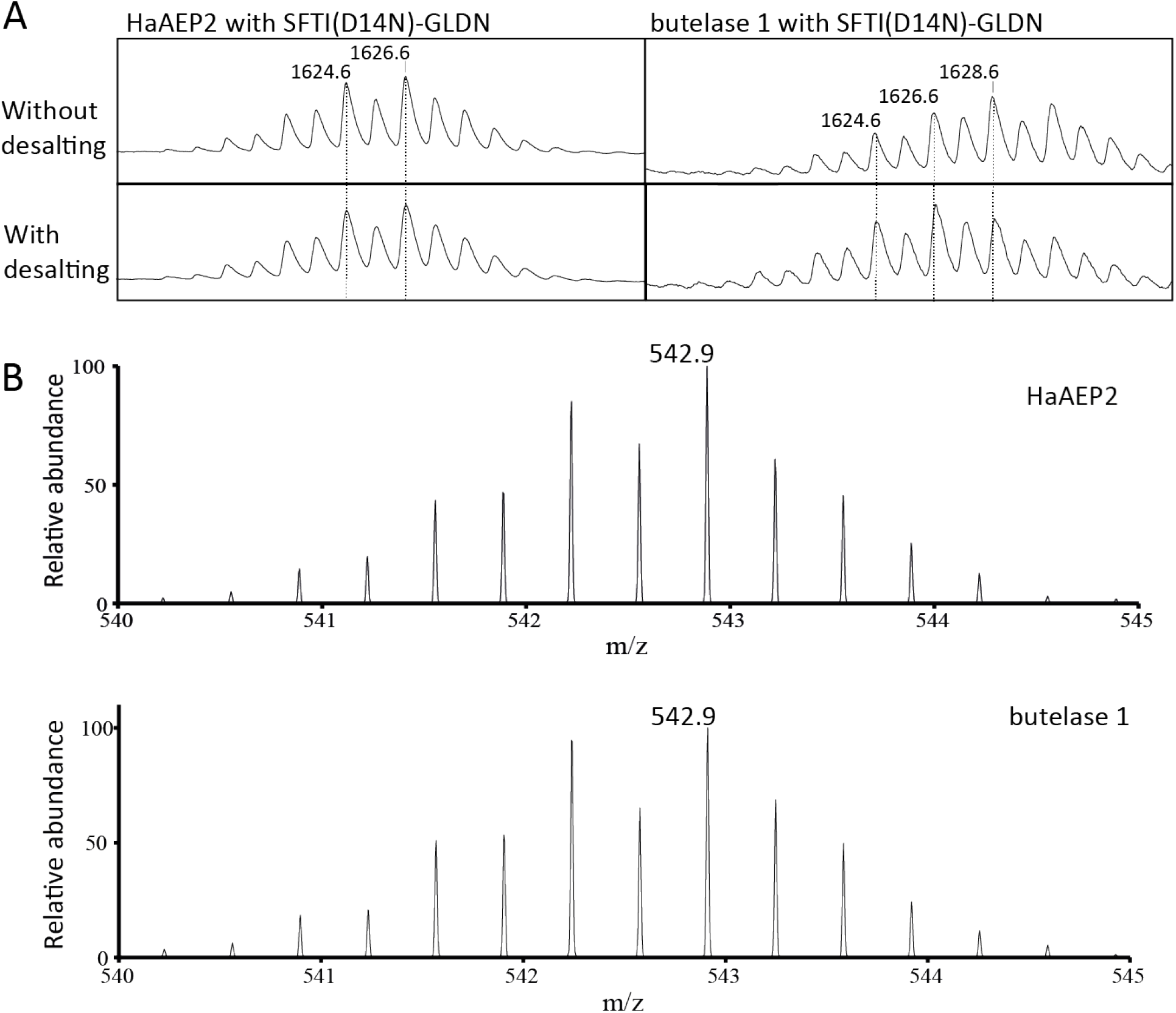
Sodium adduct testing.(**A**) Using a desalting column removes the peaks caused by the sodium adduct of cyclic SFTI(D14N) observed in MALDI-TOF MS traces with butelase 1. A sodium adduct of cyclic SFTI(D14N) resulted in a peak envelope containing both acyclic-SFTI(D14N) and the +22 sodium adduct of cyclic SFTI(D14N); however, following desalting the peak at 1628.6 corresponding to the sodium adduct is reduced. (**B**) Similarly, the expected m/z for the 3+ ion of acyclic SFTI(D14N) is observed in LC/MS due to the salt being removed during separation of the peptides by liquid chromatography resulting in the expected peak envelope.

## References

Aharoni, A., Gaidukov, L., Khersonsky, O., Gould, S.M., Roodveldt, C., and Tawfik, D.S. (2005). The ‘evolvability’ of promiscuous protein functions. Nat. Genet. 37, 73–76.

Battye, T.G.G., Kontogiannis, L., Johnson, O., Powell, H.R., and Leslie, A.G.W. (2011). iMOSFLM: a new graphical interface for diffraction-image processing with MOSFLM. Acta Crystallogr D Biol Crystallogr 67, 271–281.

Bechtold, N., Ellis, J., and Pelletier, G. (1993). *In planta Agrobacterium*-mediated gene transfer by infiltration of adult *Arabidopsis thaliana* plants. Comptes rendus de l’Académie des sciences. Série III, Sciences de la vie 316, 1194–1199.

Bernath-Levin, K., Nelson, C., Elliott, A.G., Jayasena, A.S., Millar, A.H., Craik, D.J., and Mylne, J.S. (2015). Peptide macrocyclization by a bifunctional endoprotease. Chem. Biol. 22, 571–582.

Bi, X., Yin, J., Nguyen Giang, K.T., Rao, C., Halim Nurashikin Bte, A., Hemu, X., Tam James, P., and Liu, C.-F. (2017). Enzymatic engineering of live bacterial cell surfaces using butelase 1. Angew. Chem. Int. Ed. 56, 7822–7825.

Buller, A.R., and Townsend, C.A. (2013). Intrinsic evolutionary constraints on protease structure, enzyme acylation, and the identity of the catalytic triad. Proc. Natl. Acad. Sci. U. S. A. 110, E653–661.

Chen, V.B., Arendall, W.B., III, Headd, J.J., Keedy, D.A., Immormino, R.M., Kapral, G.J., Murray, L.W., Richardson, J.S., and Richardson, D.C. (2010). MolProbity: all-atom structure validation for macromolecular crystallography. Acta Crystallogr. Sect. D Biol. Crystallogr. 66, 12–21.

Copley, S.D. (2014). An evolutionary perspective on protein moonlighting. Biochem. Soc. Trans. 42, 1684–1691.

Dall, E., Fegg, J.C., Briza, P., and Brandstetter, H. (2015). Structure and mechanism of an aspartimide-dependent peptide ligase in human legumain. Angew. Chem. Int. Ed. 54, 2917–2921.

Emsley, P., and Cowtan, K. (2004). Coot: model-building tools for molecular graphics. Acta Crystallographica Section D 60, 2126–2132.

Fisher, M.F., Zhang, J., Taylor, N.L., Howard, M.J., Berkowitz, O., Debowski, A.W., Behsaz, B., Whelan, J., Pevzner, P.A., and Mylne, J.S. (2018). A family of small, cyclic peptides buried in preproalbumin since the Eocene epoch. Plant Direct 2, 1–17.

Gruis, D., Schulze, J., and Jung, R. (2004). Storage protein accumulation in the absence of the vacuolar processing enzyme family of cysteine proteases. Plant Cell 16, 270–290.

Harris, K.S., Durek, T., Kaas, Q., Poth, A.G., Gilding, E.K., Conlan, B.F., Saska, I., Daly, N.L., van der Weerden, N.L., Craik, D.J., and Anderson, M.A. (2015). Efficient backbone cyclization of linear peptides by a recombinant asparaginyl endopeptidase. Nat. Commun. 6, 10199.

Hasegawa, H., and Holm, L. (2009). Advances and pitfalls of protein structural alignment. Curr. Opin. Struct. Biol. 19, 341–348.

Haywood, J., Schmidberger, J.W., James, A.M., Nonis, S.G., Sukhoverkov, K.V., Elias, M., Bond, C.S., and Mylne, J.S. (2018). Structural basis of ribosomal peptide macrocyclization in plants. eLife 7, e32955.

Hemu, X., Qiu, Y., Nguyen, G.K.T., and Tam, J.P. (2016). Total synthesis of circular bacteriocins by Butelase 1. J. Am. Chem. Soc. 138, 6968–6971.

Hughes, A.L. (1994). The evolution of functionally novel proteins after gene duplication. Proc Biol Sci 256, 119–124.

Jackson, M.A., Gilding, E.K., Shafee, T., Harris, K.S., Kaas, Q., Poon, S., Yap, K., Jia, H., Guarino, R., Chan, L.Y., Durek, T., Anderson, M.A., and Craik, D.J. (2018). Molecular basis for the production of cyclic peptides by plant asparaginyl endopeptidases. Nat. Commun. 9, 2411.

Jacob, F. (1977). Evolution and tinkering. Science 196, 1161–1166.

James, A.M., Haywood, J., and Mylne, J.S. (2018). Macrocyclization by asparaginyl endopeptidases. New Phytol. 2118, 923–928.

Jensen, R.A. (1976). Enzyme recruitment in evolution of new function. Annu. Rev. Microbiol. 30, 409–425.

Kaltenbach, M., and Tokuriki, N. (2014). Dynamics and constraints of enzyme evolution. J. Exp. Zool. B. Mol. Dev. Evol. 322, 468–487.

Khersonsky, O., Roodveldt, C., and Tawfik, D.S. (2006). Enzyme promiscuity: evolutionary and mechanistic aspects. Curr. Opin. Chem. Biol. 10, 498–508.

Krissinel, E., and Henrick, K. (2007). Inference of macromolecular assemblies from crystalline state. J. Mol. Biol. 372, 774–797.

Kuroyanagi, M., Yamada, K., Hatsugai, N., Kondo, M., Nishimura, M., and Hara-Nishimura, I. (2005). Vacuolar processing enzyme is essential for mycotoxin-induced cell death in *Arabidopsis thaliana*. J. Biol. Chem. 280, 32914–32920.

Lobstein, J., Emrich, C., Jeans, C., Faulkner, M., Riggs, P., and Berkmen, M. (2012). SHuffle, a novel *Escherichia coli* protein expression strain capable of correctly folding disulfide bonded proteins in its cytoplasm. Microb. Cell Fact. 11, 56.

Lu, H., Chandrasekar, B., Oeljeklaus, J., Misas-Villamil, J.C., Wang, Z., Shindo, T., Bogyo, M., Kaiser, M., and van der Hoorn, R.A. (2015). Subfamily-specific fluorescent probes for cysteine proteases display dynamic protease activities during seed germination. Plant Physiol. 168, 1462–1475.

McPhillips, T.M., McPhillips, S.E., Chiu, H.-J., Cohen, A.E., Deacon, A.M., Ellis, P.J., Garman, E., Gonzalez, A., Sauter, N.K., Phizackerley, R.P., Soltis, S.M., and Kuhn, P. (2002). Blu-Ice and the Distributed Control System: software for data acquisition and instrument control at macromolecular crystallography beamlines. Journal of Synchrotron Radiation 9, 401–406.

Mylne, J.S., and Botella, J.R. (1998). Binary vectors for sense and antisense expression of *Arabidopsis* ESTs. Plant Mol. Biol. Report. 16, 257–262.

Mylne, J.S., Colgrave, M.L., Daly, N.L., Chanson, A.H., Elliott, A.G., McCallum, E.J., Jones, A., and Craik, D.J. (2011). Albumins and their processing machinery are hijacked for cyclic peptides in sunflower. Nat. Chem. Biol. 7, 257–259.

Nguyen, G.K.T., Hemu, X., Quek, J.-P., and Tam, J.P. (2016a). Butelase-mediated macrocyclization of d-amino-acid-containing peptides. Angew. Chem. Int. Ed. 55, 12802–12806.

Nguyen, G.K.T., Wang, S., Qiu, Y., Hemu, X., Lian, Y., and Tam, J.P. (2014). Butelase 1 is an Asx-specific ligase enabling peptide macrocyclization and synthesis. Nat. Chem. Biol. 10, 732–738.

Nguyen, G.K.T., Kam, A., Loo, S., Jansson, A.E., Pan, L.X., and Tam, J.P. (2015). Butelase 1: A versatile ligase for peptide and protein macrocyclization. J. Am. Chem. Soc. 137, 15398–15401.

Nguyen, G.K.T., Qiu, Y., Cao, Y., Hemu, X., Liu, C.-F., and Tam, J.P. (2016b). Butelase-mediated cyclization and ligation of peptides and proteins. Nat. Protocols 11, 1977–1988.

Otegui, M.S., Herder, R., Schulze, J., Jung, R., and Staehelin, L.A. (2006). The proteolytic processing of seed storage proteins in *Arabidopsis* embryo cells starts in the multivesicular bodies. Plant Cell 18, 2567–2581.

Pei, J., Kim, B.H., and Grishin, N.V. (2008). PROMALS3D: a tool for multiple protein sequence and structure alignments. Nucleic Acids Res. 36, 2295–2300.

Schrodinger, L. (2010). The PyMOL Molecular Graphics System, Version 1.4.

Serra, A., Hemu, X., Nguyen, G.K.T., Nguyen, N.T.K., Sze, S.K., and Tam, J.P. (2016). A high-throughput peptidomic strategy to decipher the molecular diversity of cyclic cysteine-rich peptides. Sci. Rep. 6, 23005.

Shimada, T., Yamada, K., Kataoka, M., Nakaune, S., Koumoto, Y., Kuroyanagi, M., Tabata, S., Kato, T., Shinozaki, K., Seki, M., Kobayashi, M., Kondo, M., Nishimura, M., and Hara-Nishimura, I. (2003). Vacuolar processing enzymes are essential for proper processing of seed storage proteins in *Arabidopsis thaliana*. J. Biol. Chem. 278, 32292–32299.

Tokuriki, N., Jackson, C.J., Afriat-Jurnou, L., Wyganowski, K.T., Tang, R., and Tawfik, D.S. (2012). Diminishing returns and tradeoffs constrain the laboratory optimization of an enzyme. Nat. Commun. 3, 1257.

Winn, M. D., Ballard, C.C., Cowtan, K.D., Dodson, E.J., Emsley, P., Evans, P.R., Keegan, R.M., Krissinel, E.B., Leslie, A.G.W., McCoy, A., McNicholas, S.J., Murshudov, G.N., S. Pannu, N., Potterton, E.A., Powell, H.R., Read, R.J., Vagin, A., and Wilson, K.S. (2011). Overview of the CCP4 suite and current developments. Acta Crystallogr D Biol Crystallogr 67, 235–242.

Yang, R., Wong, Y.H., Nguyen, G.K.T., Tam, J.P., Lescar, J., and Wu, B. (2017). Engineering a catalytically efficient recombinant protein ligase. J. Am. Chem. Soc. 139, 5351–5358.

Zauner, F.B., Elsasser, B., Dall, E., Cabrele, C., and Brandstetter, H. (2018a). Structural analyses of *Arabidopsis thaliana* legumain gamma reveal the differential recognition and processing of proteolysis and ligation substrates. J. Biol. Chem. 10.1074/jbc.M117.817031.

Zauner, F.B., Dall, E., Regl, C., Grassi, L., Huber, C.G., Cabrele, C., and Brandstetter, H. (2018b). Crystal structure of plant legumain reveals a unique two-chain state with pH-dependent activity regulation. Plant Cell 30, 686–699.

Zhao, L., Hua, T., Crowley, C., Ru, H., Ni, X., Shaw, N., Jiao, L., Ding, W., Qu, L., Hung, L.-W., Huang, W., Liu, L., Ye, K., Ouyang, S., Cheng, G., and Liu, Z.-J. (2014). Structural analysis of asparaginyl endopeptidase reveals the activation mechanism and a reversible intermediate maturation stage. Cell Res. 24, 344–358.

